# Crossover formation and coordinated assembly of synaptonemal complex relies on a direct interaction between Zip1 and Zip3

**DOI:** 10.1101/2025.11.21.689840

**Authors:** Sabrina Sharmin, Karen Voelkel-Meiman, Alex J. Poppel, Amy J. MacQueen

**Author notes:** Corresponding author: Amy J. MacQueen, Wesleyan University, 238 Hall-Atwater Laboratories, 52 Lawn Avenue, Middletown CT, 06459. These authors contributed equally to the work (Sabrina Sharmin, Karen Voelkel-Meiman).

## Abstract

Several proteins collaborate to promote the crossover recombination events critical for accurate chromosome segregation during meiosis. How these “ZMM” factors (Zip2, Zip3, Zip4, Spo16, Mer3 and MutSγ) collaboratively function remains incompletely understood. We previously reported that Zip3’s abundance and activity rely on the synaptonemal complex (SC) component Zip1, and specifically on Zip1’s N-terminal residues associated with crossovers and coupling SC assembly to the crossover pathway. Here, we demonstrate that Zip3 co-immunoprecipitates Zip1 from meiotic cells independent of recombination initiation and other ZMMs, and that Zip3’s interaction with Zip1 relies on Zip1’s N terminal residues. Co-expression and pull-down experiments in bacterial cells demonstrate that Zip1 and Zip3 interact directly. Experiments to identify Zip3 regions required for the Zip1 interaction unexpectedly revealed an incorrectly annotated translational start; we also determined that Zip3’s N-terminal structured region is necessary and sufficient for the interaction, and a predicted coil downstream of Zip3’s RING domain is essential for specific activities attributed to Zip1’s N-terminal tip such as proximity labeling of Zip3 by Zip2 and the coupling of crossover recombination to SC assembly. Finally, we discovered that interaction with Zip1 protects Zip3 not only from proteasome-mediated degradation but also from post-translational modification when another ZMM is absent. We propose that direct interaction with Zip1’s N terminus orients Zip3 within a nascent ZMM ensemble in a manner that facilitates crossover formation and the coupling of crossover intermediates to SC assembly, and furthermore ensures Zip3 remains both abundant and unmodified until all requisite ZMMs have joined the group.

## Introduction

During meiosis, the maternal and paternal copies of each chromosome are successfully partitioned into separate gamete nuclei. To achieve this outcome, meiotic nuclei undergo a dramatic and unique reorganization that generates transient but stable pairwise attachments between homologous chromosomes. The formation of such chromosomal linkages in most sexually reproducing organisms relies on the meiotic DNA double-strand break (DSB) repair pathway, where programmed DNA DSBs form and are subsequently repaired in a manner that results in reciprocal exchange (crossover) events involving homologous DNA duplexes (HUNTER 2015; ZICKLER AND KLECKNER 2015). Another hallmark feature of meiotic chromosomes at this stage is a ∼100 nm wide, supramolecular multi-protein structure that spans lengthwise-aligned homologous chromosome axes along their full lengths; this structure, the synaptonemal complex (SC), provides a physical context for inter-homolog recombination intermediates (PAGE AND HAWLEY 2004; BORNER *et al*. 2023). Meiotic DNA DSB repair and SC formation reorganize DNA molecules on both local and global levels, and these widely conserved processes are executed and coordinated by specialized proteins, many of which are meiosis specific. However, the discrete molecular mechanisms that coordinate many aspects of these hallmark chromosomal events remain poorly understood.

The “ZMM” proteins (including Zip2, Zip3, Zip4, Spo16, Mer3, Msh5-Msh5) play a major role in promoting crossovers and in linking intermediate events of recombination to the initiation of SC assembly in budding yeast (Figure 1A; LYNN *et al*. 2007), and homologs of ZMMs in other organisms play similar roles (CHUA AND ROEDER 1998; NAKAGAWA AND OGAWA 1999; ZALEVSKY *et al*. 1999; AGARWAL AND ROEDER 2000; HIGGINS *et al*. 2004; JANTSCH *et al*. 2004; SNOWDEN *et al*. 2004; CHEN *et al*. 2005; FRANKLIN *et al*. 2006; TSUBOUCHI *et al*. 2006; ADELMAN AND PETRINI 2008; STRONG AND SCHIMENTI 2010; MACAISNE *et al*. 2011; CHELYSHEVA *et al*. 2012; GUIRALDELLI *et al*. 2013; REYNOLDS *et al*. 2013; DE MUYT *et al*. 2014; QIAO *et al*. 2014; RANJHA *et al*. 2014; LAKE *et al*. 2015; DUROC *et al*. 2017; DE MUYT *et al*. 2018; GUIRALDELLI *et al*. 2018; ARORA AND CORBETT 2019; LAKE *et al*. 2019). ZMM proteins co-localize as foci at presumed crossover-fated recombination sites, and their functionalities include DNA helicase activity (Mer3, Sgs1), the capacity to bind branched DNA structures (the Zip2-Zip4-Spo16 (“ZZS”) subcomplex or the Msh4-Msh5 heterodimer, aka MutSγ) and a putative SUMO E3 ligase (Zip3).

**Figure 1.**
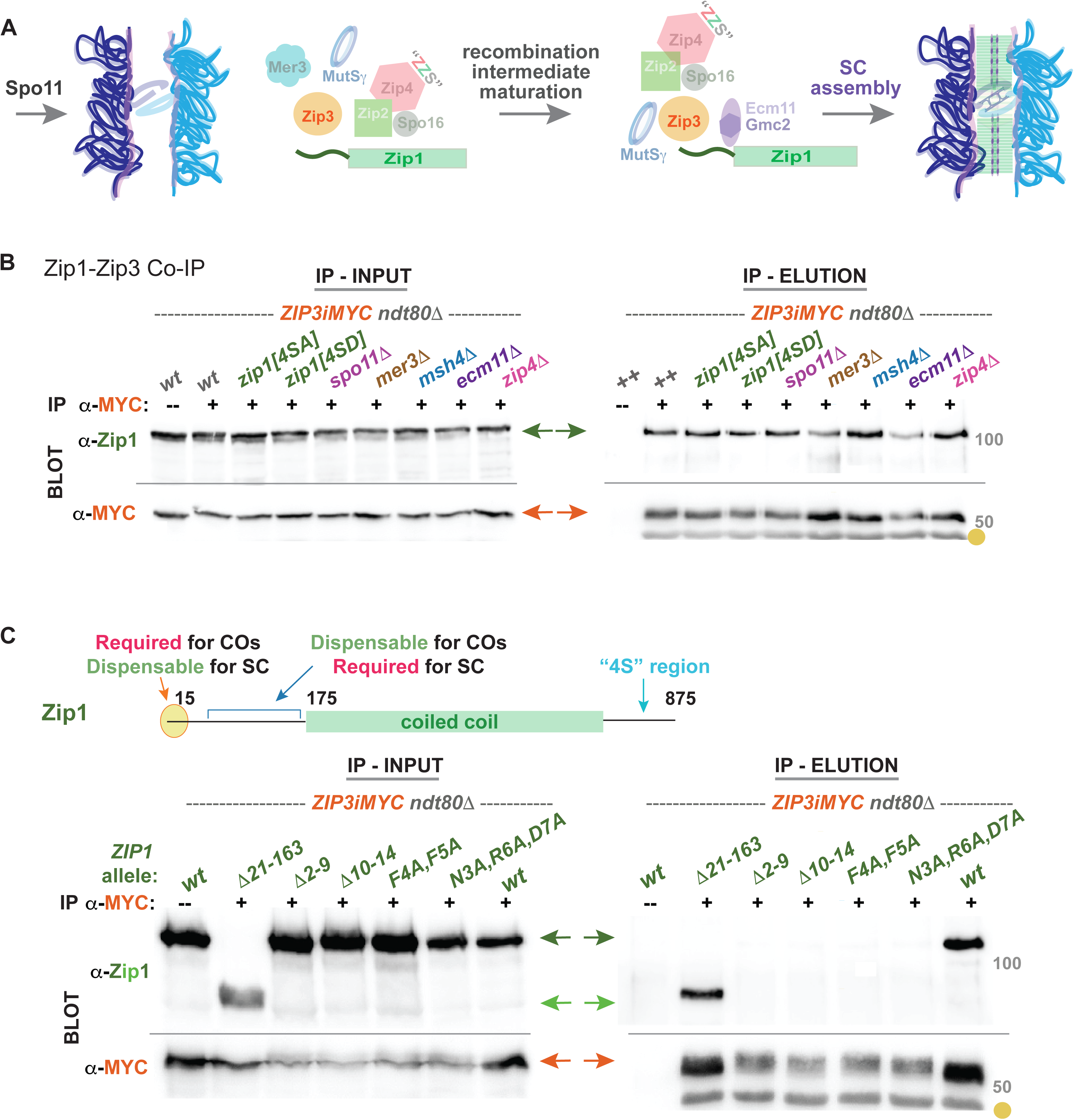
Zip3’s capacity to co-immunoprecipitate Zip1 requires Zip1’s N terminal tip but not other pro-crossover meiotic factors. Cartoon in (A) illustrates key meiotic proteins that initiate (Spo11) and facilitate ZMM-associated meiotic DNA repair events and assembly of synaptonemal complex (SC) between homologous chromosomes (shown in dark and light blue). Western blots in (B) and (C) correspond to 8% polyacrylamide gels with proteins that immunoprecipitate with Zip3iMYC from *ndt80*Δ meiotic extracts at 24 hours after introduction into sporulation conditions. Anti-MYC antibody was used to immunoprecipitate Zip3iMYC; a no-antibody control is in the first lane of each blot. Leftmost blots show input samples, corresponding to extracts prior to immunoprecipitation, while righthand blots contain elutions of Zip3iMYC from the beads and any co-immunoprecipitating proteins. Blots were probed with anti-MYC (orange arrows) or anti-Zip1 (green arrows) antibodies to visualize Zip3iMYC or Zip1. A yellow circle labels a species corresponding to the antibody. Blots in (B) show Zip3iMYC-Zip1 co-immunoprecipitation in mutants missing a key recombination or ZMM protein. *zip1-4SA and 4SD* alleles that disrupt four serines at the C terminal end of the protein, which were previously shown to disrupt recombination (CHEN *et al*. 2015). Blots in (C) show Zip3iMYC-Zip1 co-immunoprecipitation for “*zip1* tip” mutants, which disrupt the N terminal region of Zip1 and have separation-of-function phenotypes as depicted in the associated cartoon (VOELKEL-MEIMAN *et al*. 2019). Grey numbers, 50 or 100, denote protein molecular weight.

Interestingly, another factor required for ZMM-mediated crossover events in budding yeast is a structural component of the SC, Zip1. SC architecture includes two substructures that are discernable in ultrastructural images – transverse filaments are rod-like protein units that assemble in multimer fashion with their long axis perpendicular to the chromosome axis, whereas the central element substructure appears to organize transverse filament proteins at the SC’s midline. Zip1, the SC structural component with a critical role in crossover formation, dimerizes through an extensive coiled-coil region to generate the rod-like transverse filaments of the budding yeast SC (SYM *et al*. 1993; DONG AND ROEDER 2000), while Ecm11 and Gmc2 proteins form a heterocomplex that builds the SC central element (CE; HUMPHRYES *et al*. 2013; VOELKEL-MEIMAN *et al*. 2013) but these factors are dispensable for ZMM crossover recombination (VOELKEL-MEIMAN *et al*. 2016). As a central player in both major meiotic chromosomal processes, Zip1 is uniquely positioned to couple intermediate steps in recombination with the elaboration of SC structure along chromosome arms.

While many regions along the length of the Zip1 polypeptide are likely required for its capacity to serve both as a recombination protein and an SC structural component, we previously discovered that amino acids within its N-terminal tip participate exclusively in Zip1’s crossover function and are dispensable its capacity to assemble SC (VOELKEL-MEIMAN *et al*. 2019). The fact that Zip1’s N-terminal tip is one of the most highly conserved regions of the polypeptide raises the question: What molecular mechanism underpins the specialized pro-crossover role of Zip1’s N-terminus? Our prior investigation into this question revealed that Zip1’s N-tip residues appear to have a unique functional relationship with one ZMM protein, Zip3 (VOELKEL-MEIMAN *et al*. 2019). We showed that upon removal or replacement of certain N-tip residues in Zip1, cells behave as if they have lost Zip3 function with respect to crossover formation and the regulation of SUMOylated Ecm11, and that Zip1’s N-tip residues govern Zip3’s pattern of localization at polycomplex structures (aggregates of Zip1 and other SC proteins that form when these proteins are in excess). We subsequently demonstrated that these Zip1 residues are required for the proximity labeling of Zip3 by other ZMMs such as Zip2 or Msh4, and that Zip3’s abundance in the meiotic cell relies uniquely on Zip1, not any other ZMM or SC protein (VOELKEL-MEIMAN *et al*. 2024).

In aggregate, these data suggest that Zip1’s N-terminal tip residues may directly interact with Zip3. Here, we provide evidence that Zip3 interacts directly with Zip1 in a manner that relies on Zip1’s N-terminal residues, and that the two proteins may share an interaction interface at a coil adjacent to Zip3’s RING domain. We also show that, as part of or in addition to providing an activity essential to ZMM-mediated crossover formation, the Zip1-Zip3 interaction protects Zip3 from proteasome-mediated degradation and mediates a mechanism that links ZMM crossover formation to Zip3 post-translational modification.

## Methods

### Strains and crossover data

All yeast strains used in this study are isogenic to BR1919-8B (ROCKMILL AND ROEDER 1998) and are listed in Table S1. Transformation procedures to create most alleles are described in (VOELKEL-MEIMAN *^e^t al.* 202^4^). A *kanMX6-P_SCC1_* promoter cassette was inserted upstream of *PRE2* in a diploid and this was followed by a second transformation to knock out the intact *PRE2* gene to create the hemizygous meiosis deficient *PRE2* allele. A homozygous version of *kanMX6-P_SCC1_-PRE2* was created by standard genetic crossing. Diploid strains for crossover analysis are as described in (VOELKEL-MEIMAN *et al*. 2019). *zip3* alleles in this study are notated using the corrected translational start site which initiates with “MPDSI” (Figure 2), the *3xiMYC* allele has MYC inserted between amino acids 204 and 205. *Zip3[iG@20]* is a G insertion 63 nucleotides upstream, and *zip3(*Δ*A@22)* is a single A deletion 57 nucleotides upstream of the corrected translational start site.

**Figure 2.**
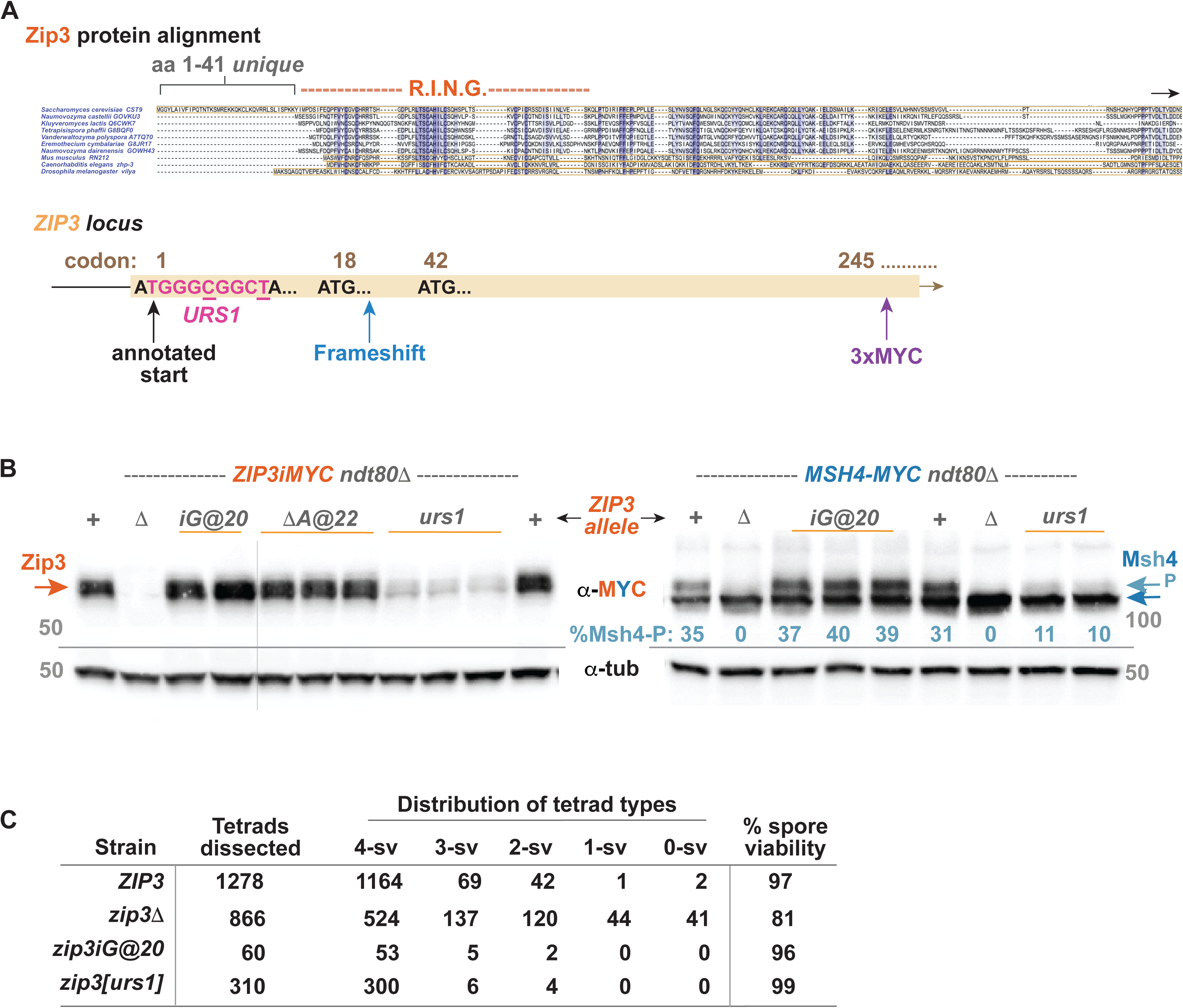
*ZIP3*’s translational start codon occurs 113 nucleotides downstream of a functional *URS1* promoter element. Upper diagram in (A) aligns the N-terminal region of *S. cerevisiae* Zip3 (top line) with Zip3 homologs from six distantly related yeast species, *Mus musculus*, *Caenorhabditis elegans*, and *Drosophila melanogaster* (performed using Uniprot (www.uniprot.org); the alignment reveals a ∼40 residue N-terminal extension unique to the *S. cerevisiae* protein. Lower cartoon illustrates the *ZIP3* gene locus as currently annotated (www.sgd.org), with the translational start codon overlapping a *URS1* promoter element (pink nucleotide sequence). The cartoon also indicates the presence of two downstream *ATG* codons in the same reading frame (at codon positions 18 and 42), and the approximate position of frameshift allele disruptions (insertion of a G just after codon 20 or deletion of an A just before codon 23), as well as the position of the internal 3xMYC epitope utilized to monitor Zip3 abundance in this study. Finally, the underlined nucleotides within the *URS1* sequence (pink) correspond to those simultaneously changed to create the *zip3[urs1]* allele. (B) Proteins extracted from mid-meiotic prophase arrested (*ndt80*Δ) cells homozygous for *ZIP3iMYC* were separated on an 8% polyacrylamide gel and transferred to nitrocellulose, which was then sequentially probed with anti-MYC antibody and anti-tubulin as a loading control. Left blot contains meiotic extracts from cells homozygous for *zip3iMYC* alleles encoding a frameshift (*iG@20* or Δ*A@22*) or carrying a disrupted *URS1* sequence (*TGGGtGGCa*). Quantitation of Zip3iMYC levels can be found in Supplemental Figure S2 and Supplemental File S7. Right blot corresponds to strains carrying the same *zip3* alleles but without an internal MYC tag, instead these strains are homozygous for *MSH4-13xMYC*, which enables us to detect phosphorylated Msh4 protein (lighter blue arrow at right), whose presence is associated with successful *ZMM* crossover formation (HE *et al*. 2020; VOELKEL-MEIMAN *et al*. 2024). Light blue numbers on right blot gives the % of total Msh4-MYC that corresponds to the phosphorylated species. Grey numbers at left and right (50 or 100) refer to molecular weight (MW). Table in (C) gives spore viability information for strains homozygous for *zip3* null, frameshift or *urs1* alleles. For each strain, the tetrad products of meiosis were dissected and evaluated for whether all four (4-sv), three (3-sv), two (2-sv), one (1-sv) or no (0-sv) spores show viability (grow into colonies). Percent spore viability (last column) is the total number of observed viable spores over the number expected if all four spores of every dissected tetrad is viable, multiplied by 100.

### Co-immunoprecipitation

55 ml of sporulating cultures were collected at a 24-hour timepoint. Cells were resuspended in freshly made lysis buffer (50mM Tris pH8, 200mM NaCl, 10mM EDTA, 1x protease inhibitor, 1mM PMSF, 10% glycerol, 1mM DTT, 0.5% IgePAL CA-630) and all subsequent steps were carried out at 4°C. Cells were lysed using ∼50% volume of 5 mm glass beads with vortexing at top speed (20 seconds on/10 seconds rest for 20 cycles). 50 µl of Protein G magnetic beads (Invitrogen, #10003D) and 2 µl mouse anti-MYC (Invitrogen 9E10) antibody were prepared as per manufacturer’s recommendation using protein LoBind tubes (Eppendorf). 50 µl of each input sample was collected prior to lysate incubation with antibody-tagged protein G beads and mixed with 50 µl of protein sample buffer (100 mM Tris pH 6.8, 20% glycerol, 4% SDS, 200mM DTT, 0.2% bromophenol blue) supplemented with 30mM DTT. After a 90-minute incubation, the beads were washed three times with 1xPBST and eluted in 30 µl sample buffer. Samples were heated for 10 minutes at 65°C, then centrifuged for 5 minutes at 13,000 rpm.

### Bortezomib drug treatment

Bortezomib was solubilized in DMSO according to the manufacturer’s protocol (Thermo Fisher Scientific; catalog number J60378.MA). Diploid strains were grown overnight in YPADU media to an OD of 0.4-0.5, cells were pelleted, washed and resuspended in sporulating media (2% KAc) at a 1:5 ratio and split into 4 flasks. At 18-hours post-meiotic induction, DMSO or bortezomib were added to the flasks to make the final bortezomib concentration 0µM, 125µM, 250µM and 500µM. After a 6-hour incubation period with the drug, protein pellets were isolated and processed from 5ml cultures via TCA precipitation (see below).

### Western blot and streptavidin blotting

Protein was extracted from 5ml sporulating culture via TCA precipitation and processed as described in (VOELKEL-MEIMAN *et al*. 2024) and incubated overnight at 4°C with primary antibody in blocking buffer for Invitrogen Mouse anti-MYC antibody, or 1xPBST for all other antibodies: Mouse anti-MYC (Invitrogen 9E10) was used at 1:5000 for Zip3-iMYC and at 1:10,000 for all other MYC detection, rabbit anti-Zip1 and rat anti-tubulin YOL1/34 (Abcam) was used at 1:10,000; mouse anti-ubiquitin at 1:2000 and rabbit anti-Fpr3-C (gift from Dr. Jeremy Thorner) at 1:50,000. HRP-conjugated secondary antibodies - goat anti-mouse (Jackson ImmunoResearch), donkey anti-rabbit, and goat anti-rat (Santa Cruz) - were used at 1:10,000 dilution in 1xPBST for 1 hour at room temperature. ECL Prime western blotting detection reagent (Amersham) was used to detect protein on the membrane. A Syngene G:Box was used for signal detection using the Syngene GeneTools software. Blots were processed and analyzed using ImageJ (https://imagej.net/ij/index.html).

### Cytological analysis and imaging

Meiotic chromosome nuclei were surface spread and imaged as in (VOELKEL-MEIMAN *et al*. 2016). The following primary antibodies were used: affinity purified rabbit anti Zip1 (YenZym Antibodies, LLC, 1:200), mouse anti-Gmc2 (raised against purified Gmc2, ProSci Inc., 1:800), rabbit anti-Red1 (gift from G.S. Roeder.1:200) and guinea pig anti-Gmc2_Ecm11 (raised against a co-purified protein complex; ProSci Inc., 1:800). Alexa Fluor dye-conjugated secondary antibodies were obtained from Jackson ImmunoResearch and used at 1:200 dilution.

### Recombinant protein expression and amylose affinity pulldown assay

*ZIP1* residues 1-348 were cloned into pMAT11, providing an N-terminal 6xHisMBP-Zip1 polypeptide (cleavable using tobacco etch virus (TEV) protease), a kind gift from Owen Davies (University of Edinburgh). *ZIP1* mutations were introduced using QuikChange XL site directed mutagenesis. The *ZIP3* residues 1-152, and all altered versions of *ZIP3* were cloned into pRSFDuet (Novagen) creating a 6xHIS tag-Zip3 polypeptide by One Step SLIC-cloning. All plasmids were confirmed by plasmid sequencing. Constructs were co-expressed in Rosetta 2 (DE3) cells (Novagen) in 2xYT media, expression induced at an OD_600_ of 0.7 with 0.5 mM isopropyl-β-D-thiogalactosidase (IPTG) for 16 hours at 25°C. Cells were lysed by sonication in 20 mM Tris pH 8.0, 300 mM KCl and 1µM PMSF, centrifuged to remove cell debris at 20,000x*g* for 20 minutes. Lysates were added to an amylose resin column equilibrated with 20 mM Tris pH 8.0, 300 mM KCl, 2mM DTT and incubated at 4°C for 1 hour with rocking. Supernatant was collected, amylose washed with 20 mM Tris pH 8.0, 300 mM KCl, 2mM DTT, and elution of the MBP bound complexes was accomplished using wash buffer plus 30mM D-maltose and collected as 1 ml samples. PAGE analysis was done to examine protein using 4x Laemmli buffer and proteins were visualized with coomassie stain.

## Results

### Zip3 co-immunoprecipitates Zip1 independent of Spo11 and ZMMs but relies on N-terminal residues of Zip1 required for ZMM crossover formation

We used co-immunoprecipitation (co-IP) to gain additional insight into the physical and functional relationship between Zip1’s N-terminal residues and Zip3 during meiosis. Zip1 was previously reported to elute with immunoprecipitated Zip3 from *rad50S* meiotic cells (AGARWAL AND ROEDER 2000). *rad50S* mutants initiate meiotic recombination but the Spo11-mediated DNA DSBs fail to undergo early processing (ALANI *et al*. 1990). The fact that Zip1 and Zip3 co-IP from *rad50S* cells indicates that some form of a Zip1-Zip3 ensemble forms independent of early interhomolog recombination intermediates. To ask whether the Zip1-Zip3 interaction observed by co-IP depends on meiotic DSBs, or requires other ZMMs or SC components, we evaluated Zip3’s ability to co-IP Zip1 in mutants missing Spo11, Mer3, Zip4, Msh4 or Ecm11.

Anti-MYC was used to immunoprecipitate Zip3iMYC (a version of Zip3 with 3 copies of the MYC epitope located at an internal position within the polypeptide) from *ndt80* extracts at the 24-hour timepoint; most cells of the BR genetic background at this timepoint contain mature interhomolog recombination intermediates (DNA joint molecules) and full-length SC structures (VOELKEL-MEIMAN *et al*. 2012; VOELKEL-MEIMAN *et al*. 2016), owing to the absence of the Ndt80 transcription factor required for progression beyond late prophase (XU *et al*. 1995). We detected both Zip3iMYC and Zip1 in the elution fraction from this sample, but not in elutions from control immunoprecipitations lacking the MYC antibody (Figure 1B). The capacity of Zip3iMYC to co-IP Zip1 from cells at late prophase aligns with our prior observation that Zip3’s abundance depends upon Zip1 at this stage (VOELKEL-MEIMAN *et al*. 2024). While most immunoprecipitation experiments in this study were performed at this 24-hour timepoint, we also determined that Zip3iMYC can detectably co-IP Zip1 at earlier timepoints (21- and 17-hour; Supplemental Figure S1). Zip1 was not detected in Zip3iMYC eluates at 13-hours but was hardly detected in the “input” sample at this timepoint, suggesting Zip1 abundance is below the limit of detection of our antibody at this early meiotic stage.

These co-IP data indicate that Zip3 indirectly or directly interacts with Zip1 throughout meiotic prophase, including at late pachytene when ZMM-associated recombination intermediates and SC structures are most abundant. We next asked whether other meiotic factors are important for the Zip1-Zip3 interaction. We first examined *spo11* mutants as ZMM factors may interact with one another differently depending on the presence of recombination intermediates. For example, we previously showed that proximity labeling of Zip3 by Zip2 occurs independent of Spo11 but that proximity labeling of Zip3 by Msh4 relies on Spo11 and Mer3 activity (VOELKEL-MEIMAN *et al*. 2024). Here we found that Zip3iMYC readily co-IPs Zip1 from meiotic cells missing Spo11, indicating that Zip1-Zip3 ensembles exist independent of recombination initiation.

Next, we asked about the dependency relationship between the Zip1-Zip3 ensemble and other ZMM factors or components of the SC. The Zip2, Zip4, and Spo16 proteins form a biochemically stable subcomplex (“ZZS”; DE MUYT *et al*. 2018), and Zip2’s capacity to proximity label Zip3 depends upon its ZZS partners (VOELKEL-MEIMAN *et al*. 2024). However, we found that Zip3iMYC readily co-IPs Zip1 even in the absence of Zip4, indicating that ZZS is not essential for the formation of a Zip1-Zip3 ensemble. We furthermore found the Zip1-Zip3 interaction does not require the Mer3 DNA helicase, the MutSγ component, Msh4, nor another core SC structural component, Ecm11 (Figure 1B). Taken together, our results indicate that Zip3 interacts with Zip1 independent of most, if not all, other known ZMM proteins and SC components.

We previously showed that Zip1’s N-terminal tip residues (amino acids 1-15) play a critical role in mediating ZMM-mediated crossovers and that these residues enable Zip2 to proximity label Zip3 (VOELKEL-MEIMAN *et al*. 2024). If the mechanistic basis of Zip1’s N-tip crossover activity is an interaction with Zip3, these Zip1 tip residues should be required for Zip3’s capacity to co-IP Zip1. We found this to be the case: Zip3iMYC failed to co-IP any of four altered versions of Zip1 (encoded by *zip1* alleles *[Δ2-9], [Δ10-14], [F4A,F5A], [N3A,R6A,D7A])* that alter N-tip residues and diminish or abolish Zip1’s ZMM crossover activity (VOELKEL-MEIMAN *et al*. 2019; Figure 1C). On the other hand, we found Zip3iMYC co-IPs the Zip1[Δ21-163] protein, which contains an in-frame deletion in a downstream region of Zip1’s N terminus that renders the protein SC assembly-deficient but crossover-proficient (VOELKEL-MEIMAN *et al*. 2016; Figure 1C).

Not all crossover-deficient *zip1* mutants abolish the Zip3-Zip1 co-IP. The *zip1* “4SA” mutant (carrying four serine replacements toward the C terminus of the protein) was previously reported to be deficient for interhomolog crossovers and SC assembly owing to an inability to be phosphorylated at these residues by the Mek1 kinase (CHEN *et al*. 2015). The Zip1[4SA] and Zip1[4SD] proteins are both readily co-immunoprecipitated with Zip3iMYC protein from meiotic extracts (Figure 1B). These findings in aggregate thus strengthen a functional link between i) the crossover-promoting activity specifically attributed to Zip1’s N terminal tip and ii) a physical interaction with the meiosis-specific E3 ligase, Zip3.

### Structure-function analysis of Zip3 reveals an incorrectly annotated translational start and that Zip3’s N terminal region is necessary and sufficient for interaction with Zip1

To learn about molecular features of the Zip3 protein important for its interaction with Zip1’s N terminal tip, we created several *zip3* mutant alleles that encode alterations in distinct regions of Zip3’s polypeptide chain. Alignment of *S. cerevisiae* Zip3’s polypeptide sequence with the homologs from several other yeast, as well as mouse, *Drosophila*, and *C. elegans*, highlights conserved regions throughout the N-terminally located RING domain and a downstream coil (Figure 2A), but less conservation or predicted structure in the C terminal half of the protein. The alignment also highlights an N terminal ∼40 amino acid extension that is unique to *S. cerevisiae* Zip3. Curiously, residue 42 of *S. cerevisiae* Zip3 is a methionine. Furthermore, a *URS1* transcriptional regulatory element (*TGGGCGGCTA*) overlaps the first four annotated codons of *S. cerevisiae ZIP3,* raising the possibility that this region of the *ORF* may have promoter function (Figure 2A). These observations led us to question the annotated translational start for *S. cerevisiae ZIP3*.

To ask whether *ZIP3*’s true translational start is codon 42 (the second of two methionine codons downstream of the *ATG* embedded in the *URS1* element) we generated and analyzed putative frameshift alleles. We created *zip3* alleles with either a single nucleotide insertion (*iG@20*) or deletion (Δ*A@22*), positioned immediately after codon 20 or before 23 respectively, which is upstream of only one *ATG* codon (codon 42) prior to an essential histidine residue (H80; CHENG *et al*. 2006; SERRENTINO *et al*. 2013). We created these mutations in the context of *ZIP3iMYC* (carrying an in-frame 3xMYC epitope) and a non-epitope tagged *ZIP3* gene. If either of the two *ATG* codons upstream of codon 42 is utilized as the translational start for *ZIP3*, no Zip3iMYC protein should be detectable in protein extracts and no functional Zip3 protein should be produced in either of these frameshift alleles. In contrast to this expectation, we observed ample Zip3iMYC protein in meiotic cells homozygous for the *iG@20* or *ΔA@22* mutation (Figure 2B). Furthermore, *zip3[iG@20]* strains exhibit normal Msh4 phosphorylation at late meiotic prophase (Figure 2B; Msh4 phosphorylation is associated with ZMM crossover activity and requires Zip3 function) and wild-type spore viability (Figure 2C). These data indicate that meiotic Zip3 function relies on a protein that starts with the RING domain region of the polypeptide, better matching the N termini of homologous Zip3 proteins (Figure 2A). We will use our updated annotation for Zip3, with the N-terminus consisting of amino acids “MPDSI”, throughout the remainder of this study.

We also gathered data suggesting the nine base pair *URS1* sequence overlapping the previously annotated translational start is important for *ZIP3* expression. We created a mutant allele (*zip3[urs1])* that changes two nucleotides (cytosine 5 to thymine and thymine 9 to adenine) shown to be important for *URS1* function during either meiosis (cytosine 5) or mitosis (thymine 9) (GAILUS-DURNER *et al*. 1997). Although the phenotypes of frameshift alleles (above) indicate this nucleotide sequence is not part of *ZIP3*’s coding region, we designed the double mutation to compromise *URS1* activity while preserving potential protein-coding capacity. Consistent with a defect in *URS1* function, strains homozygous for an epitope-tagged version of the *zip3[urs1]* allele produce much less Zip3iMYC protein (∼25% of a wild type allele; Figure 2B and Supplemental Figure S2A). The most interesting aspect of this mutant is the very low level of Zip3 in the *zip3[urs1]* strain is sufficient for full *ZIP3* function: First, spore viability and crossover recombination across seven genetic intervals on chromosomes *III* and *VIII* in *zip3[urs1]* resemble wild type rather than the *zip3* null (Figure 2C, Table 1). Furthermore, while the *zip3* null exhibits a severe deficiency in SC assembly and a high frequency of nuclei displaying polycomplex (an assembly of SC and recombination proteins that is characteristic of cells with defects in SC initiation or elaboration) *zip3[urs1]* meiotic cells display robust SC assembly and a very low frequency of polycomplex (Supplemental Figure S2B). Finally, the abundance of SUMOylated forms of Ecm11 is increased ∼1.5-2-fold when Zip3 function is missing (Supplemental Figure S2C), but *zip3[urs1]* meiotic cells show wild-type levels of SUMOylated Ecm11. Thus, a relatively low level of Zip3 (barely detectable using our blotting conditions) is sufficient to provide full Zip3 function to meiotic cells.

**Table 1.**
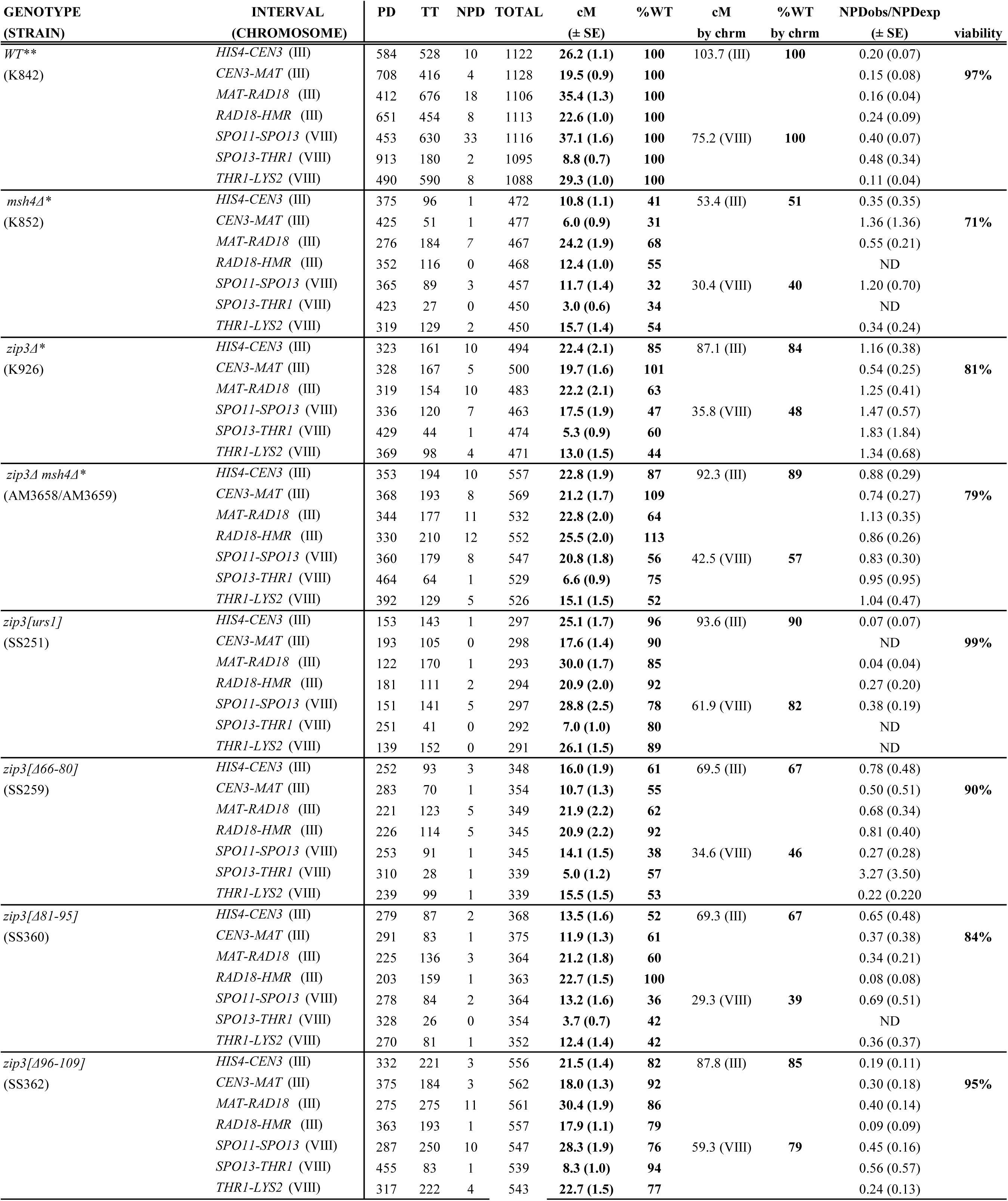
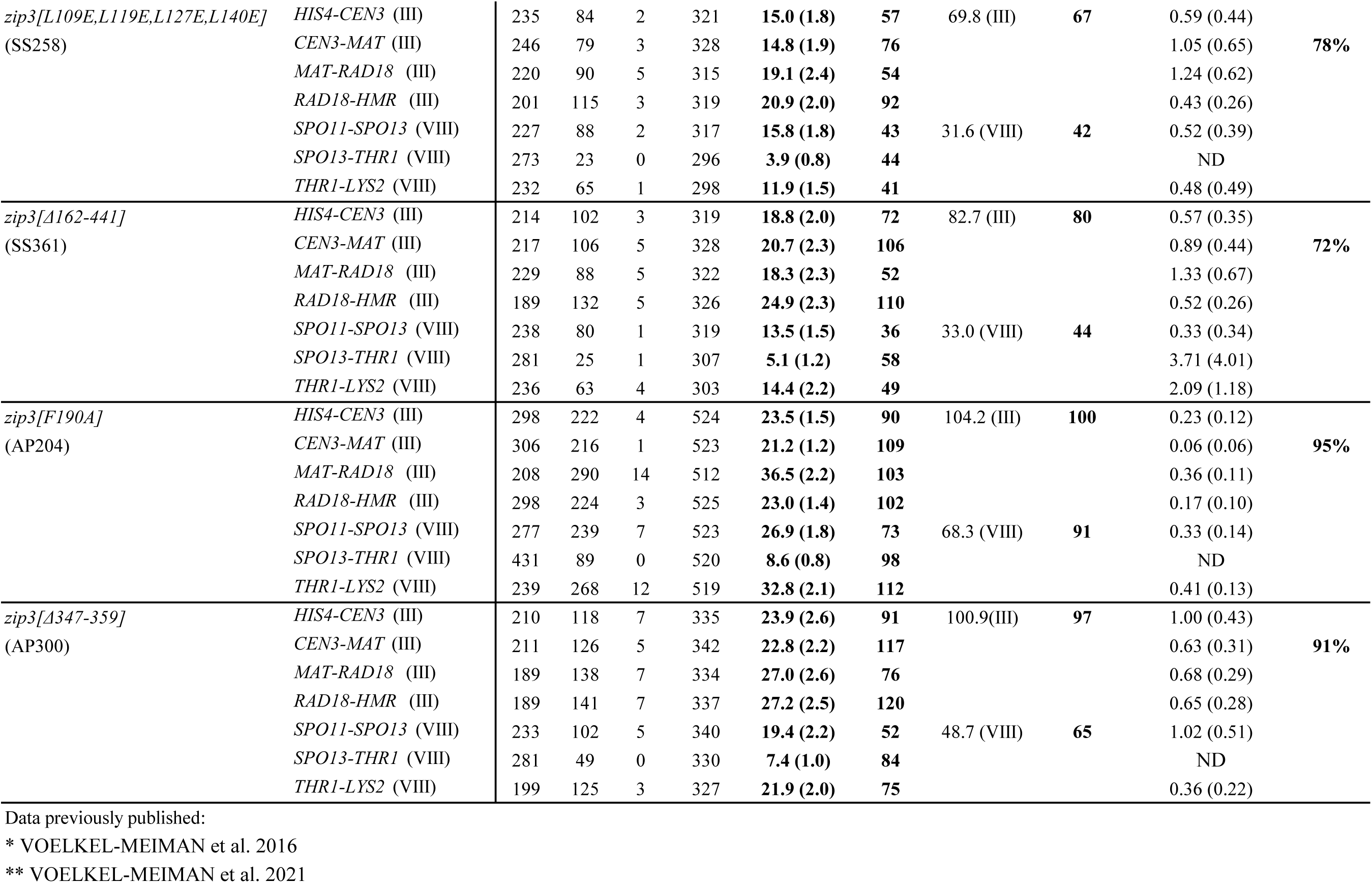
Crossover frequency in *zip3* alleles Genetic linkage analysis to measure crossover levels in *zip3* strains. Map distances and genetic interference values were calculated using tetrad analysis and coefficient of coincidence measurements as described previously (VOELKEL-MEIMAN *et al*. 2013; VOELKEL-MEIMAN *et al*. 2015; VOELKEL-MEIMAN *et al*. 2019). Table gives map distances (standard errors) and their corresponding percentages of the wild-type values for individual intervals, and for the entire chromosome (by summing the intervals on III or VIII). For intervals marked (ND), interference measurements are not obtainable using the coefficient of coincidence method due to an absence of NPD tetrads. Crossover frequencies for strains marked with an * or ** have been previously published (VOELKEL-MEIMAN *et al*. 2016; VOELKEL-MEIMAN *et al*. 2022, respectively).

We next evaluated a series of in-frame deletion and point mutants to identify residues within the Zip3 polypeptide important for Zip3’s capacity to co-IP Zip1 (alleles illustrated in Figure 3A). Remarkably, we found through this structure-function analysis that the C terminal two-thirds of Zip3 (corresponding to residues 162-441 and predicted to be predominantly unstructured), is dispensable for the Zip1-Zip3 interaction in cells: A 3xMYC-tagged version of Zip3[Δ162-441] readily co-IPs Zip1 from meiotic extracts (Figure 3B). Consistently, Zip3[F190A]iMYC and Zip3[Δ347-359]iMYC proteins, which carry alterations within small regions of conservation in the C terminal half of the protein, also co-IP Zip1 from meiotic cells.

**Figure 3.**
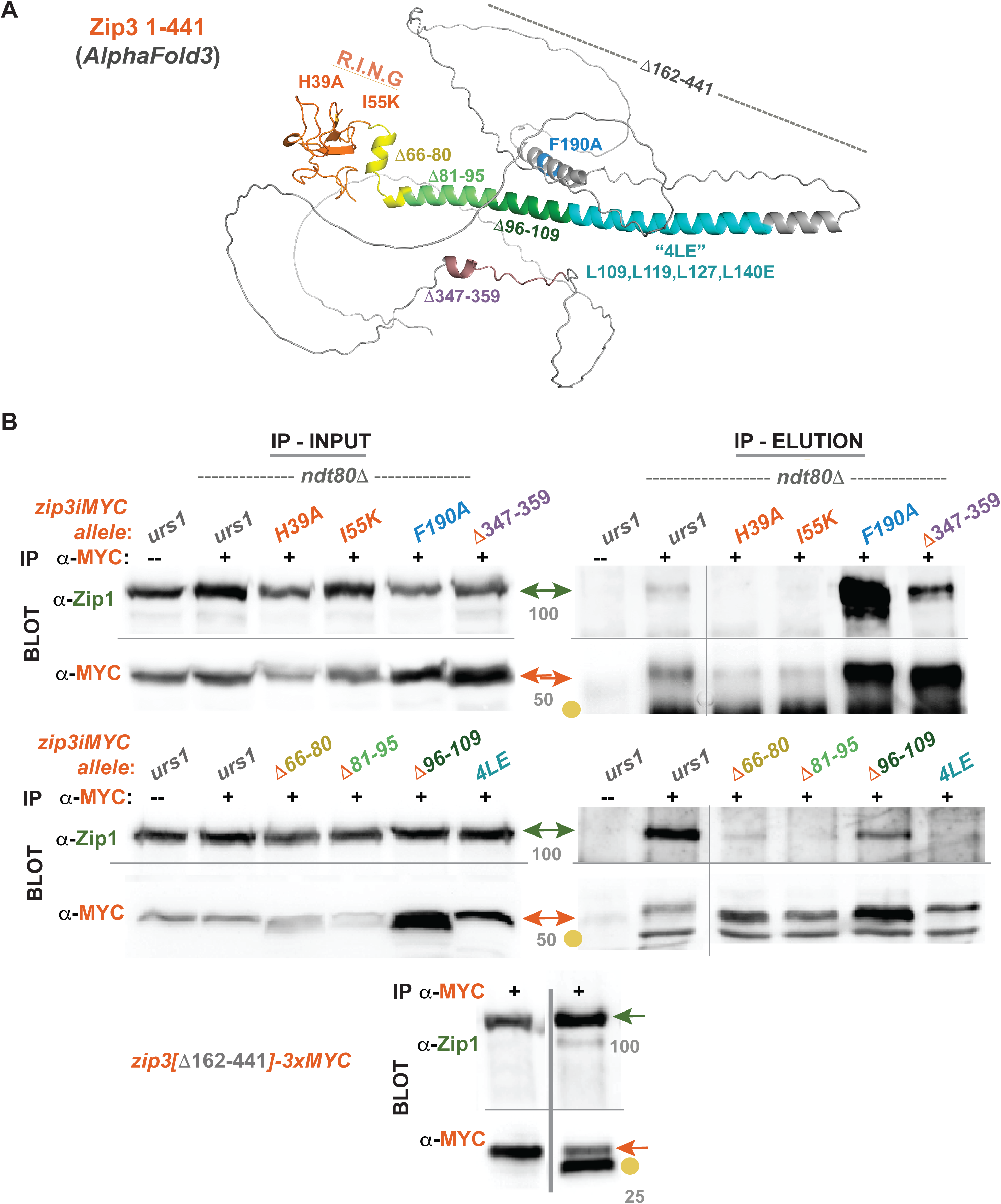
Zip3 N terminal residues are required for co-immunoprecipitation of Zip1. Upper diagram in (A) shows a molecular structure proposed by *AlphaFold3* for the Zip3 polypeptide with updated annotation (meaning Zip3’s N-terminus begins “MPDSI”). Labeled and distinctly colored regions (rendered in Pymol) indicate disruptions in particular *zip3* mutant alleles generated for this study. Western blots in (B) correspond to Zip3-Zip1 co-immunoprecipitation experiments: Meiotic extracts from *ndt80*Δ cells were incubated with anti-MYC antibody to immunoprecipitate Zip3iMYC (a no-antibody control is in the first lane of each blot). Left-most blots (and to the left of grey bar in the lowest blot) contain input samples (prior immunoprecipitation), while blots on the right contain Zip3iMYC elutions and any co-immunoprecipitating proteins. Anti-MYC (orange arrows) or anti-Zip1 (green arrows) was used to detect Zip3iMYC or Zip1, respectively, on blots (8% polyacrylamide gels were used in all except the lowest blot, which is a 12% gel). A yellow circle labels a species that corresponds to the anti-MYC antibody. Strains in these experiments carry various *zip3* alleles, as indicated at top of blots, allele colors correspond to regions indicated in (A). Since many of the *zip3* mutants have low Zip3 abundance (input blots in B and Supplemental Figure S4), *zip3[urs1]iMYC* is used as a control comparison (Zip3 abundance is low (Supplemental Figure S2A) but Zip1 is nevertheless detected as a co-immunoprecipitating protein in the *zip3[urs1]-iMYC* strain). Grey numbers correspond to molecular weight (MW).

By contrast, most alterations affecting residues in N terminal third of Zip3 compromise Zip3’s capacity to co-IP Zip1. Mutations affecting Zip3’s RING domain (H39 and I55K; CHENG *et al*. 2006; SERRENTINO *et al*. 2013) abolish Zip3’s ability to co-IP Zip1 (Figure 3B), although the fact that Zip3iMYC protein itself is barely detectable in eluates from *H39A* and *I55K* strains raises the caveat that the negative Zip1 result stems from a technical limitation, such as insufficient Zip3 levels or compromised antibody-Zip3iMYC interaction, rather than a failed Zip1-Zip3 interaction.

Like Zip3[H39A] and Zip3[I55K], four additional altered forms of Zip3iMYC containing disruptions to a predicted coil downstream of the RING domain also showed low abundance relative to non-mutant Zip3iMYC (Figure 3B “inputs”), but in these four alleles (*zip3[Δ66-80], zip3[Δ81-95], zip3[Δ96-109],* and *“4LE”*= *zip3[L109E,L119E,L127E,L140E])* the level of Zip3 protein resembles that from strains homozygous for the *urs1* allele where Zip1 is detectably co-immunoprecipitated. Thus, we carried out Zip3iMYC-Zip1 co-IP experiments on these coil-disrupting *zip3* alleles in parallel to a *zip3*[*urs1]* mutant. Interestingly, Zip1 was severely diminished but detectable in eluates from Zip3[Δ66-80] and Zip3[Δ96-109] immunoprecipitations but absent from immunoprecipitated eluates of Zip3[Δ81-95] or the Zip3[4LE] protein. Our findings indicate that the Zip1-Zip3 interaction, as measured by co-IP from meiotic cells, relies on residues within a predicted coil downstream of Zip3’s RING domain (in addition to crossover-associated residues at the N terminal tip of Zip1) and does not require the unstructured C terminal region that comprises more than half of the Zip3 protein.

### Zip1 and Zip3 interact directly, through their N terminal regions

To investigate whether the interaction between Zip1 and Zip3 is direct, we co-expressed MBP-tagged Zip1 and His-tagged Zip3 partial proteins in *E. coli* and determined if the two proteins co-elute from amylose resin (Figure 4A). We found that upon co-expression, Zip3’s N-terminal structured region (amino acids 1-152) appeared in the fractions of MBP-tagged Zip1-N (amino acids 1-348) eluted from amylose, but not in the control elutions of free MBP from amylose (Figure 4B). These data demonstrate a direct interaction between the N terminal regions of Zip1 and Zip3.

**Figure 4.**
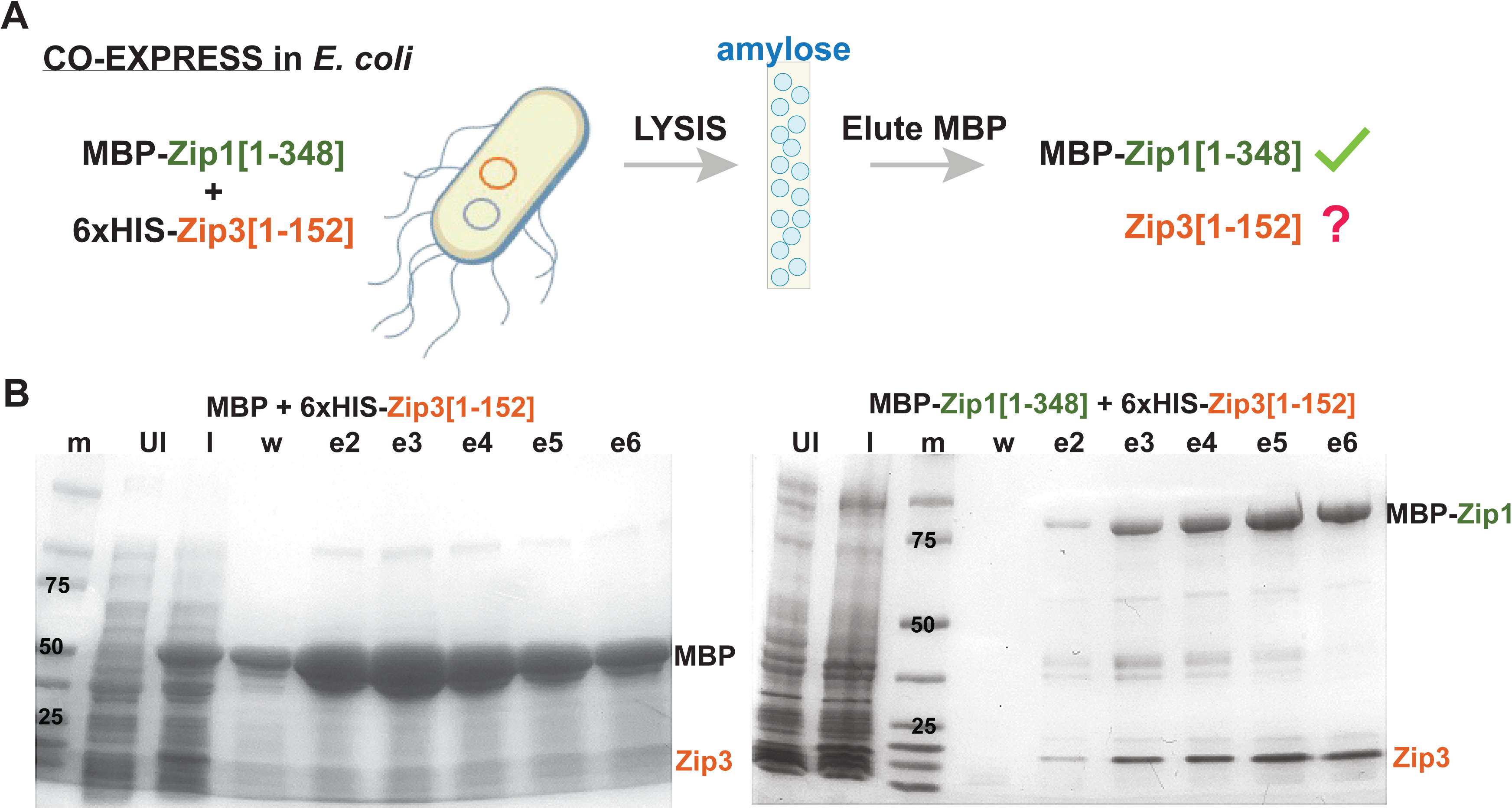
Zip3 co-purifies with MBP-Zip1 from bacterial extracts. Cartoon in (A) illustrates experimental design: Two expression plasmids, one encoding MBP (maltose binding protein) fused to an N-terminal fragment of Zip1 (amino acids 1-348) and one encoding a Zip3 fragment (residues 1-152) with an N terminal six-histidine residue tag, were introduced into *E. coli* and the expression of these proteins stimulated using IPTG. A strain carrying only an MBP-encoding plasmid in addition to the 6xHIS-Zip3[1-152] expression plasmid served as a control. MBP or MBP-Zip1 was purified from control and experimental *E. coli* cell lysates using amylose beads, and proteins in consecutive 1 mL eluates (e2-e6) were examined on a protein gel stained with Coomassie. Left gel in (B) corresponds to the MBP + 6xHIS-Zip3[1-152] control and the right gel corresponds to the experimental samples. Samples from uninduced (UI) and induced (I) whole cell extracts and the last wash (w) are also included on the gel. Labels at right indicate position of MBP-Zip1, MBP, and Zip3 species. Protein molecular weight marker sizes are indicated in the (m) lane.

Does the Zip1-Zip3 *in vitro* interaction depend on Zip1 or Zip3 residues previously implicated in the interaction? Consistent with our co-IP results from meiotic cells (Figure 1C), the Zip1[F4A, F5A] or Zip1[Δ10-14] alterations, both of which abolish MutSγ crossovers (VOELKEL-MEIMAN *et al*. 2019), destroy MBP-Zip1[1-348]’s capacity to pull down Zip3[1-152] from bacterial extracts (Supplemental Figure S3). On the other hand, the Zip1[N3A, R6A, D7A] disruption (which reduces but may not abolish ZMM crossovers VOELKEL-MEIMAN *et al*. 2019), and the Zip1[Δ21-163] disruption (which prevents SC assembly but not Zip1’s ability to promote MutSγ crossovers nor Zip3’s ability to co-immunoprecipitate Zip1 (VOELKEL-MEIMAN *et al*. 2015; VOELKEL-MEIMAN *et al*. 2019; Figure 1C)) each preserve at least some of Zip1[1-348]’s capacity to pull down Zip3[1-152] in the co-expression experiment, although the *in vitro* interaction may be weakened in these mutant contexts (Supplemental Figure S3). Amylose pulldown data thus largely agree with the results of our meiotic cell co-immunoprecipitation and indicate that the structured N-terminal region of Zip3 mediates a direct physical interaction with Zip1, with Zip1’s N-tip, crossover-associated residues playing an essential role in the interaction.

We furthermore explored the ability for MBP-Zip1[1-348] to pull down various altered forms of Zip3[1-152], but with less conclusive results. Together with MBP-Zip1[1-348], we co-expressed six altered forms of Zip3[1-152]: H39A, I55K, Δ66-80, Δ81-95, Δ96-109, and “4LE”:L109E, L119E, L127E, L140E. None of the six Zip3 partial proteins eluted with MBP-Zip1[1-348] from amylose (Supplemental Figure S4), however we cannot rule out the possibility that one or more of these disruptions to Zip3 cause an intrinsically unstable protein whose non-specific aggregation precludes an interaction with MBP-Zip1. For all constructs apart from Zip3[“4LE”], the induced Zip3 protein was readily detected in whole cell extracts, but even the positive control shows little Zip3[1-152] protein in the soluble supernatant after disrupting cells by sonication (see “sup” versus “pellet” lanes in control blot, Supplemental Figure S3).

### Interaction with Zip1 protects Zip3 from proteasome-mediated degradation

We previously reported that Zip3’s cellular abundance plummets upon Zip1 removal, suggesting that Zip3’s stability depends upon an interaction with Zip1 (VOELKEL-MEIMAN *et al*. 2024). Consistent with this possibility, disruptions to Zip3 that diminish its capacity to immunoprecipitate Zip1 also render the protein less abundant in cells (see above). We thus quantified total Zip3 abundance in *ndt80*-arrested meiotic yeast cells homozygous for each of our nine *zip3* alleles (Supplemental Figure S5A). We found that while Zip3 devoid of its entire C terminal half is highly abundant, disruptions to its RING or downstream coil region causes a severe diminishment in Zip3 levels. Alterations to residues within Zip3’s RING (H39A and I55K) and the two internal deletions closest to the RING rendered Zip3 the least abundant (∼10% of the wild type level), while the distal deletion (Δ96-109) or the Zip3[4LE] disruption show a less severe but substantial reduction in Zip3 that is reminiscent of Zip3’s abundance in cells missing Zip1 (∼20-30% of the wild type).

We reasoned that if Zip3[Δ96-109] or Zip3[4LE] protein instability is due solely to the loss of an interaction with Zip1, Zip1 removal from these cells should result in no further diminishment in Zip3 abundance; we found this to be the case particularly for Zip3[4LE], while Zip3[Δ96-109] abundance appeared mildly diminished by loss of Zip1 (Supplemental Figure S5B). Interestingly, the robust abundance of Zip3[F190A] and Zip3[Δ162-441] proteins is also less sensitive (F190A) or completely insensitive ([Δ162-441) to the presence of Zip1 (Supplemental Figure S5B). Unlike Zip3[4LE] or Zip3[Δ96-109] however, these C-terminally disrupted Zip3 proteins maintain a capacity to co-IP Zip1 (Figure 3). From these data, we conclude the two halves of Zip3 have opposing roles in Zip3’s stability in the meiotic cell: Zip3’s N terminal structured region promotes stability likely through mediating an interaction with Zip1. On the other hand, Zip3’s C terminal half is dispensable for stability and instead appears to facilitate Zip3 degradation when Zip1 is absent.

We explored whether interaction with Zip1 protects Zip3 from proteasome-mediated degradation using both a pharmacological and a genetic approach. We first interfered with proteasome function by adding bortezomib to sporulating cultures for approximately six hours immediately prior to the 24-hour timepoint when cells were collected and evaluated for protein abundance. Consistent with compromised proteasome function, we observed an increase in ubiquitinated protein species in extracts from cells exposed to bortezomib relative to control (untreated) cultures (Figure S6A). We observed that *zip1* cells exposed to 500 µM bortezomib display substantially increased Zip3 levels relative to control *zip1* cells exposed only to DMSO (Figure 5A), indicating that the proteasome is responsible for at least some of the Zip3 instability triggered by loss of Zip1. We note that concentrations of bortezomib at 500 µM and above show reduced Zip3 and Zip1 abundance (see Figure S6B for 500 µM data), suggesting this level of drug negatively impacts progression through meiosis.

**Figure 5.**
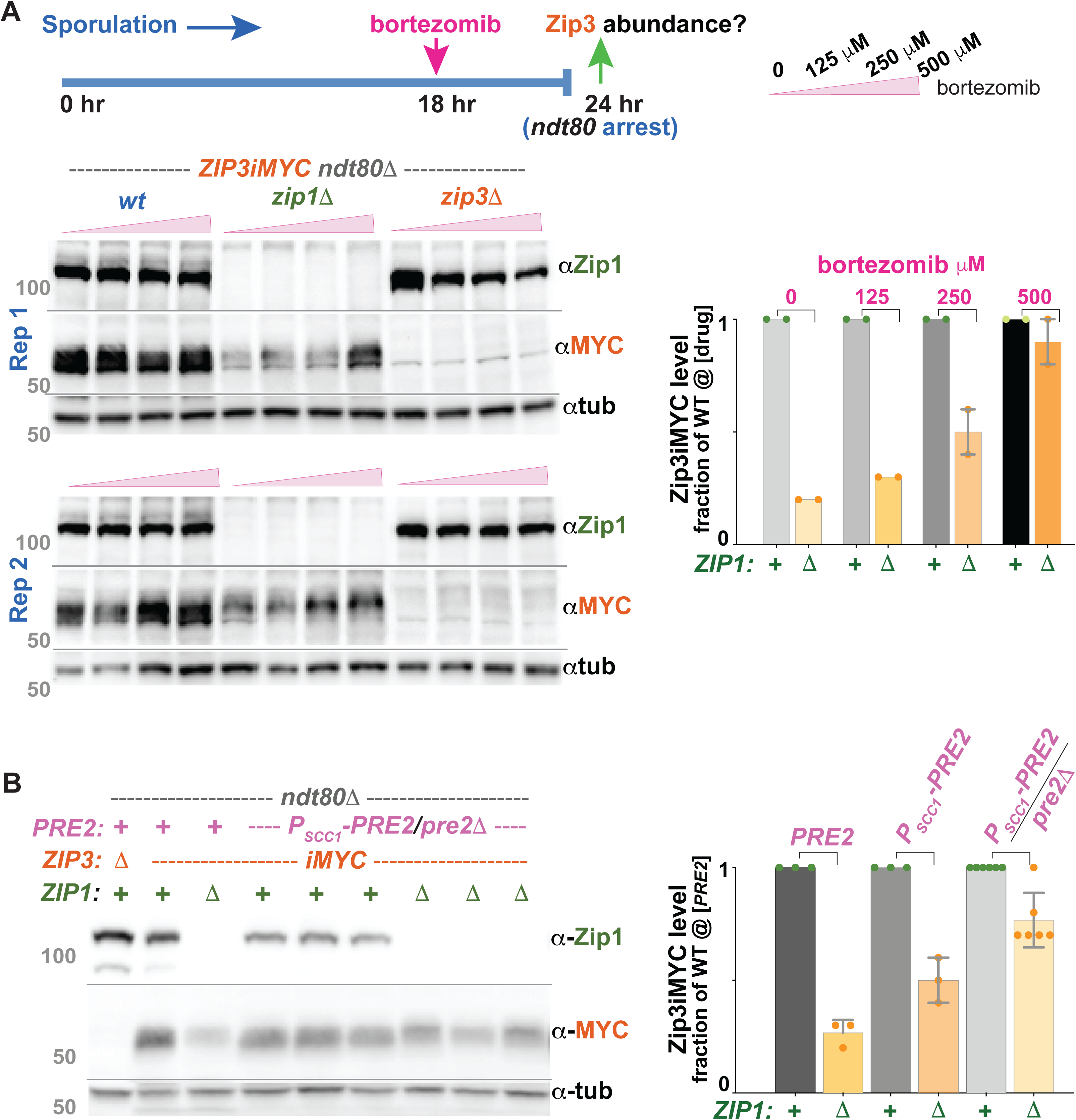
Zip1 protects Zip3 from proteasome-mediated degradation. Cartoon in (A) illustrates the experimental design for pharmacological diminishment of proteasome function in meiotic cells. Cultures of *ndt80* cells are exposed to bortezomib 18 hours after introduction to sporulation conditions and harvested six hours later to create extracts. Meiotic cell extracts were run on 8% polyacrylamide gels and examined for Zip1 and Zip3 protein abundance by western blot (lower panel); tubulin level was used as a loading control (lower blot). Grey numbers correspond to molecular weight (MW). Graph at right plots Zip3iMYC level relative to the *ZIP1*+ strain at a given bortezomib dose, which is set to 1; presence or absence of Zip1 indicated below *x* axis). Colored circles in graph indicate individual values, shaded bar indicates the mean, and error bars indicate the range. (B) Proteasome function is manipulated genetically in *ZIP1+* and *zip1Δ* strains by placing the *PRE2* gene under transcriptional control of a vegetative-specific promoter, *P_SCC1_.* Blot shows Zip3iMYC levels in meiotic extracts, examined on an 8% polyacrylamide gel. The first three lanes carry *PRE2+* controls while the remaining lanes contain extracts from strains trans-heterozygous for *P_SCC1_-PRE2* and a *pre2Δ* null allele. Three biological replicates of a *ZIP1+* version of this *P_SCC1_-PRE2/pre2Δ* strain are followed by three biological replicates of a *zip1Δ* version of the strain. Anti-Zip1, anti-MYC, and anti-tubulin antibodies were sequentially used as probes to evaluate the abundance of each protein. Grey numbers correspond to molecular weight (MW). Graph at right indicates Zip3iMYC levels in the *zip1Δ* relative to the *ZIP1+* version of a given *PRE2* genotype. Also included are data from strains homozygous for *P_SCC1_-PRE2* (blots provided in Supplemental File S7, along with all raw data plotted in graphs). Colored circles indicate individual values, shaded bar indicates mean, and error bars indicate standard deviation. We note that the presence of bortezomib in the culture, or genetic reduction in *PRE2* activity, each negatively impacts total Zip1 and Zip3 protein levels (see Supplemental Figure S6B).

We next interfered with proteasome function using a genetic approach, by creating a meiotic “shut-off” allele of *PRE2*, which encodes a core proteasome component (BAUMEISTER *et al*. 1998). This was done by placing *PRE2* under the transcriptional control of the *SCC1* promoter, which is not expressed well during meiosis (KLEIN *et al*. 1999). We observed a greater abundance of Zip3 in *zip1* cells hemizygous or homozygous for *P_SCC1_*-*PRE2*, relative to *zip1* cells with full proteasome function (Figure 5B), consistent with the outcome of pharmacological proteasome inhibition discussed above. Taken together, our findings indicate that Zip1 protects at least a substantial fraction of Zip3 from proteasome-mediated degradation during meiosis.

### Zip3’s Zip1-interacting residues promote crossovers and proximity to Zip2

The N-terminal tip residues of Zip1 (amino acids 1-15) specifically facilitate MutSγ crossover formation and are not required for Zip1’s capacity to assemble SC (VOELKEL-MEIMAN *et al*. 2019). We thus anticipated that Zip3 residues essential for interacting with Zip1’s tip would also be essential for crossover formation. To address this question, we i) evaluated Msh4 phosphorylation at late meiotic prophase (a molecular marker that correlates with successful MutSγ crossover formation (HE 2018; VOELKEL-MEIMAN *et al*. 2024) and ii) monitored crossover frequency directly, using genetic linkage analysis across eight intervals on chromosomes *III* and *VIII* (Table 1).

We found that Msh4 phosphorylation is undetectable in the late meiotic prophase cells of *zip3[I55K], zip3[Δ81-95], zip3[4LE],* or *zip3[Δ162-441]* mutants, and is substantially reduced in cells homozygous for *zip3[Δ66-80]*, *zip3[Δ96-109],* or *zip3[Δ347-359]* but remains at wild-type level in the *zip3[F190A]* mutant (Figure 6A). As expected, the Msh4 phosphorylation phenotype of each *zip3* mutant aligns well with crossover recombination frequency (Table 1). We note that as reported previously (VOELKEL-MEIMAN *et al*. 2019), the *zip3* null displays a milder crossover deficiency on chromosome *III* relative to chromosome *VIII*, and relative to the chromosome *III* crossover deficiency of a *msh4* null. While this result might imply that some MutSγ-mediated crossovers exist on chromosome *III* when Zip3 is absent, this is not the case: The *zip3 msh4* double mutant displays the same high level of crossovers on *III* as the *zip3* null (Table 1 and VOELKEL-MEIMAN *et al*. 2019). We interpret these data to mean that Zip3 prevents the formation or resolution of a subset of recombination intermediates (at least on chromosome *III*) formed by a non-ZMM mechanism (both in the *msh4* mutant and in otherwise wild-type cells). However, because chromosome *III* crossover frequencies are less informative, we focus on crossover frequencies across chromosome *VIII*, where the *zip3* and *msh4* null phenotypes closely resemble one another. Here, crossover levels track well with Msh4 phosphorylation: *zip3[Δ81-95], zip3[4LE],* and *zip3[Δ162-441]* mutants display a null-like crossover phenotype while crossovers are moderately reduced in the *zip3[Δ96-109]* or *zip3[Δ347-359]* mutant and close to wild-type level in *zip3[F190A]* (Table 1). The *zip3[Δ66-80]* mutant departs somewhat from the correlation, as Msh4 phosphorylation in this mutant is mildly reduced like *zip3[Δ96-109]* (Figure 6A), but its crossover phenotype is more severe and resembles the *zip3* null (Table 1). Taken together, these data suggest that the N-terminal regions of Zip1 and Zip3 involved in a direct interaction play a critical role in ZMM crossover formation. The mutant data also indicate that (like Zip1) more C terminal regions of Zip3 also provide an essential pro-crossover role.

**Figure 6.**
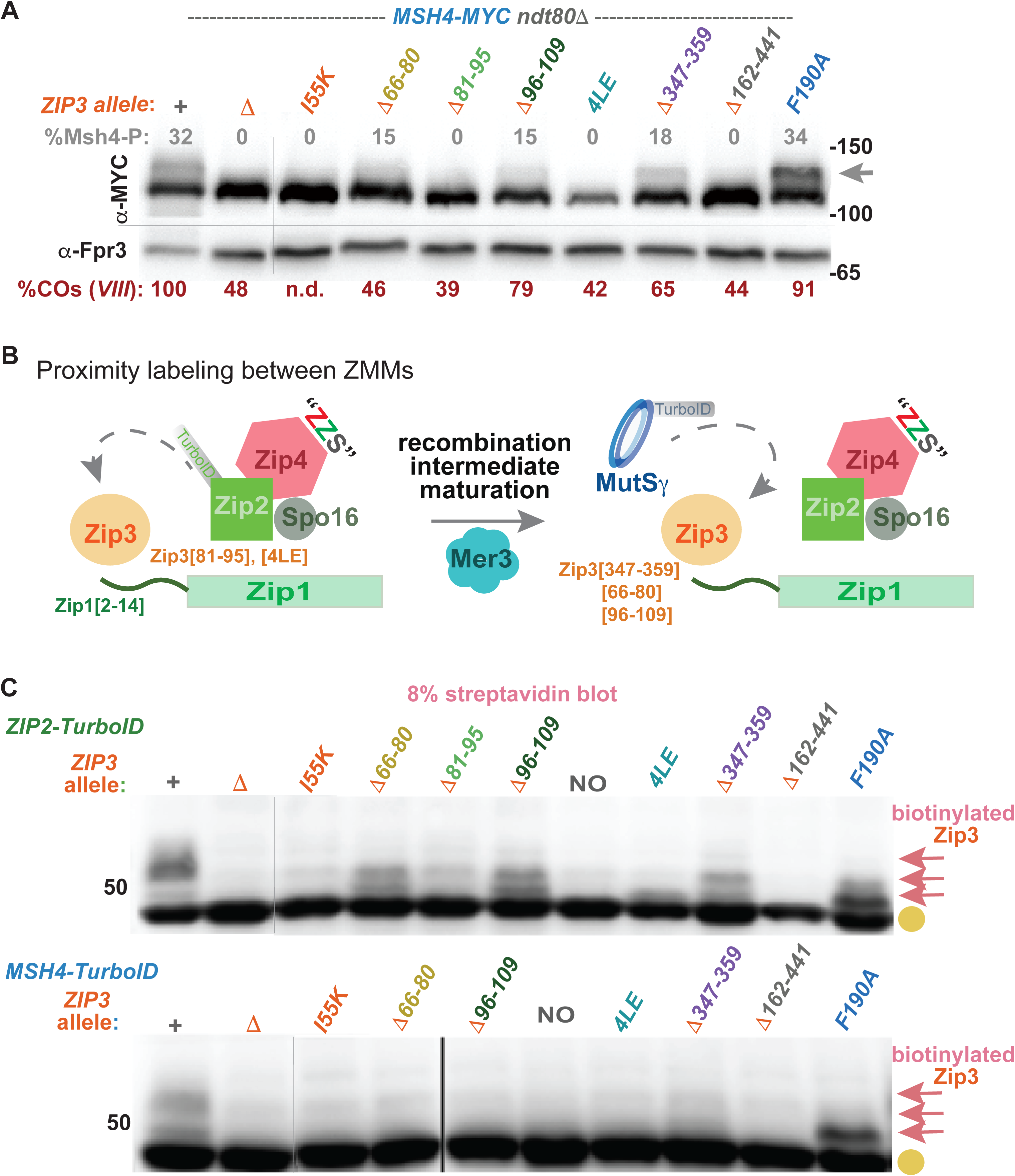
Crossover-associated events such as Msh4 phosphorylation and proximity labeling by Zip2- and Msh4-TurboID require multiple regions across Zip3, including a RING-adjacent coil. Western blot in (A) shows Msh4-MYC protein from *ndt80Δ* strains homozygous for *zip3* alleles; cells were harvested after 24 hours in sporulation media. Phosphorylated Msh4-MYC appears as a slower migrating species in 8% polyacrylamide gels, indicated by arrow. Percentage of total Msh4-MYC that corresponds to the phosphorylated version is indicated above each lane (grey); values are an average of two biological replicates; see Supplemental File S7 for raw data. Fpr3 (below horizontal line) served as a loading control. % of wild type crossover frequency across chromosome *VIII* for each strain is given in red below the blot (Table 1). Numbers at right indicate molecular weight. Cartoon in (B) illustrates how interaction with Zip1’s N terminus might orient Zip3 within an early ZMM ensemble nearby to Zip2, such that Zip2-TurboID protein can proximity label Zip3. Based on prior data (VOELKEL-MEIMAN *et al*. 2024), we envision that proximity of Zip3 to MutSγ (and particularly the Msh4 component of MutSγ) occurs only after a discrete maturation step of the recombination intermediate involving the Mer3 helicase. Blots in (C) display biotinylated proteins extracted from *ndt80Δ* cells homozygous for *ZIP2-TurboID* (green) or *MSH4-TurboID* (blue) and separated on an 8% polyacrylamide gel. *TurboID* strains are homozygous for *zip3* point mutations or in-frame deletion alleles (listed at top of blots). Gold circles indicate naturally biotinylated proteins, pink arrows indicate biotinylated Zip3 proteins. Shown are representative blots; two biological replicates were examined for all strains. Note that no interpretation can be made about the capacity for Zip3[Δ*162–441]* to be proximity labeled by either Zip2 or Msh4-TurboID; while no truncated protein was visible on the streptavidin blot at the predicted molecular weight position (using both 8% and 12% gels), we cannot rule out a technical issue preventing transfer of this small protein to the membrane.

Proximity labeling analysis using fusions between ZMMs and the yeast-optimized biotinylase enzyme, TurboID (BRANON *et al*. 2018; LAROCHELLE *et al*. 2019) furthermore suggest that Zip3’s N-terminal interaction with Zip1 may facilitate its proper positioning within the ZMM ensemble. Proximity labeling of Zip3 by Zip2-TurboID (as assessed on a streptavidin blot) is not dependent on recombination initiation, but is strongly disrupted by removal of Zip4 or Spo16, and by N terminal tip alterations to Zip1 that prevent Zip1’s proper interaction with Zip3 (VOELKEL-MEIMAN *et al*. 2024). These observations led to the proposal that a core recombinosome ensemble containing Zip2, Zip4, Spo16, Zip1, and Zip3 can assemble independent of DNA recombination intermediates and that Zip1’s N terminal tip is important for positioning Zip3 at these recombination ensembles (Figure 6B). Similarly, here we show that strains homozygous for alterations to Zip3 that abolish a detectable interaction with Zip1 in co-IP experiments - such as *zip3[I55K] zip3[Δ81-95],* and *Zip3[4LE]* - show a dramatic reduction in the proximity labeling of Zip3 by Zip2-TurboID (Figure 6C). These data support the possibility that a direct interaction between Zip1 and Zip3 serves to position Zip3 within an early recombinosome ensemble in a manner that brings Zip3 close to Zip2 (Figure 6B).

On the other hand, the Zip3[Δ66-80] and Zip3[Δ96-109] proteins, which are crossover-defective but still weakly interact with Zip1 in co-IP experiments, and the crossover-defective Zip3[Δ347-359] protein that robustly co-IPs Zip1, each appears to be proximity labeled by Zip2-TurboID, albeit possibly to a reduced extent (Figure 6C). It is interesting that this class of *zip3* mutants (defined not by an abolished Zip1-Zip3 interaction but instead by a reduced level of Zip3 proximity labeling by Zip2 and diminished crossovers) has another distinguishing feature - an intermediate level of Msh4 phosphorylation (Figure 6A). The observed diminishment in proximity labeling of Zip3 by Zip2 in this group of mutants may reflect an inability to position Zip3 properly within the early recombinosome ensemble or may be due to disabled Zip3 function that prevents the formation of a later recombinosome ensemble where Zip2 possibly for a second time comes into proximity of Zip3 (Figure 6B).

By contrast with the Zip2-Zip3 proximity interaction which is recombination independent, proximity labeling of Zip3 by Msh4-TurboID depends not only on the core ZZS factors and Zip1 but also on proteins involved in the enzymology of recombination such as the Spo11 endonuclease and Mer3 DNA helicase (Figure 6B; VOELKEL-MEIMAN *et al*. 2024), which indicates that the Msh4-Zip3 proximity interaction depends on successful completion of intermediate steps in the recombination pathway. Consistent with this idea, we observe that Msh4-TurboID fails to biotinylate Zip3 in every one of our *zip3* mutant alleles that confers a crossover defect; only wild-type Zip3 and Zip3[F190A] proteins are detectably biotinylated by Msh4-TurboID (Figure 6C).

We failed to observe biotinylated Zip3[Δ162-441] protein in Zip2-TurboID or Msh4-TurboID strains using either 8% or 12% gels, but we cannot rule out the possibility that the biotinylated form of Zip3[Δ162-441] fails to transfer to the membrane in the context of our experimental conditions. Finally, we note that Zip3 proteins biotinylated by Zip2-TurboID in our *zip3* mutants show a shifted position on the blot relative to biotinylated Zip3 in the wild type strain due to a direct or indirect defect in Zip3 post-translational modification (see below).

### Zip3’s RING-adjacent coil ensures SC assembly is linked to recombination while residues throughout its length prevent accumulation of hyperSUMOylated Ecm11

In addition to promoting MutSγ crossover formation, Zip3 is required for SC assembly initiating from recombination sites (TSUBOUCHI *et al*. 2008) and also somehow limits the accumulation of SUMOylated forms of the SC structural component, Ecm11 (HUMPHRYES *et al*. 2013). To identify regions and residues of Zip3 involved in these SC-associated functions, we evaluated our nine *zip3* mutants for their capacity to assemble SC and to control abundance of SUMOylated forms of Ecm11.

Like Zip2 and Zip4, Zip3 is essential for SC assembly from recombination sites. However, unlike these other ZMMs, Zip3 is dispensable for SC formation at centromeres and instead inhibits SC assembly from centromeres when signals linked to recombination are absent (TSUBOUCHI *et al*. 2008). Thus, while SC is completely absent from *zip2* and *zip4* null mutants at late meiotic prophase (CHUA AND ROEDER 1998; TSUBOUCHI *et al*. 2006), a limited number of SC structures assemble from centromere sites in at least a fraction of nuclei from *zip3* null mutants. In *zip3* cells arrested at late prophase owing to the *ndt80* mutation, some SC structures achieve substantial lengths (VOELKEL-MEIMAN *et al*. 2019; Figure 7A, B).

**Figure 7.**
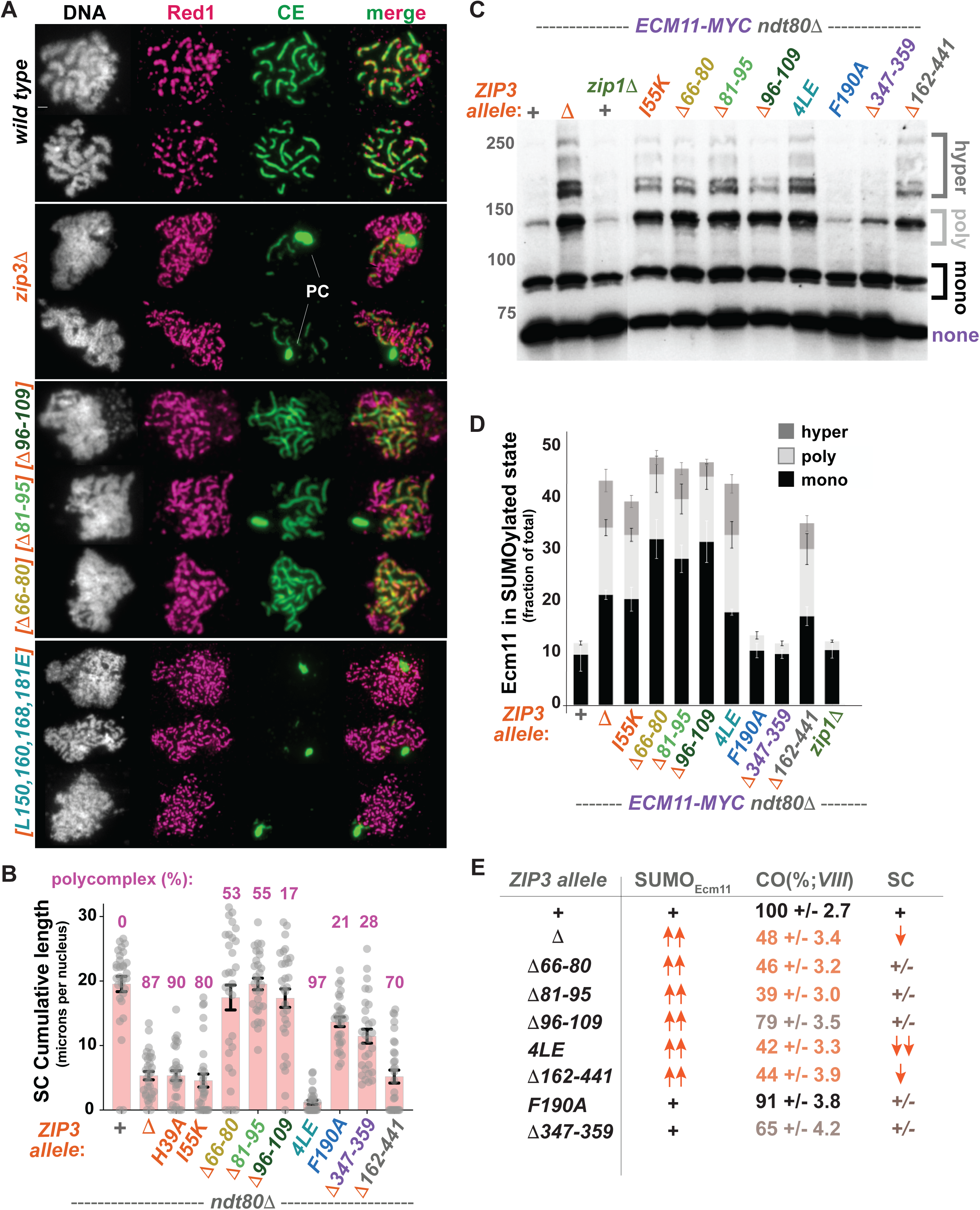
The capacity to attenuate Ecm11 SUMOylation is not strictly correlated with crossover proficiency nor SC assembly in *zip3* non-null mutants. Images in (A) show surface-spread chromosomes from strains homozygous for *ndt80Δ* and either *ZIP3+* (top two rows), a *zip3* null allele (second two rows), the *zip3[Δ96-109]*, *zip3[Δ81-95]*, *zip3[Δ66-80]* allele (next three rows, respectively), or the *zip3[L150E, L160E, L168E, L181E]* allele (last three rows). The Red1 chromosome axis protein is shown in magenta, while SC central element proteins are labeled green; DAPI labels DNA (white). Large aggregates of SC-containing proteins (polycomplex, PC) are indicated in the *zip3* images. Bar, 1 micron. Plot in (B) gives the total cumulative length of SC (microns per nucleus; each grey circle corresponds to a value for a single nucleus), with a shaded bar indicating mean and error bars indicating standard deviation. The percentage of nuclei examined with a polycomplex is given at top of graph. Western blot in (C) shows Ecm11-MYC protein from *ndt80Δ* strains homozygous for indicated alleles. Extracts were prepared after 24 hours of sporulation, when a small fraction of Ecm11 can be found in either mono, poly, or hyperSUMOylated forms, detected as slower-migrating species on the blot. Cells lacking Zip3 function show an enrichment for hyperSUMOylated Ecm11. Numbers at left indicate molecular weight. Graph in (D) plots the fraction of total Ecm11 protein in a mono, poly, or hyperSUMOylated form for each strain; data plotted are from three independent experiments and error bars representing standard deviation. Summary table in (E) lists the Ecm11-SUMO, chromosome *VIII* crossover, and SC assembly phenotypes of wild type and mutant *zip3* alleles alongside one another. Raw data for plots are supplied in Supplemental File S7.

Our nine *zip3* alleles could be classified into three phenotypic groups with respect to SC assembly in *ndt80* late meiotic prophase arrested cells. Alleles disrupting Zip3’s RING domain or removing the C terminus (*zip3[H39A]*, *zip3[I55K]*, and *zip3[Δ162-441]*) exhibit a null phenotype, with a reduced number of SC structures on surface-spread meiotic chromosomes and a large fraction of these nuclei displaying a substantial aggregate of SC proteins called a polycomplex (Figure 7A, B). In a second phenotypic class represented by *zip3[4LE]*, SC assembly is nearly abolished (Figure 7A, B): Polycomplexes were observed in almost every meiotic nucleus from *zip3[4LE]* cells, but linear SC structures were detected less often relative to the *zip3* null. This interesting phenotype suggests the Zip3[4LE] protein may be incapable of triggering SC assembly from recombination sites but proficient at inhibiting SC assembly from centromeres (TSUBOUCHI *et al*. 2008). Finally, a third phenotypic class - represented by the three internal deletions disrupting an extended coil adjacent to Zip3’s RING domain - exhibit cumulative lengths of SC nearly as extensive as the wild type, in addition to polycomplex structures (Figure 7A, B). Thus, Zip3’s RING-adjacent coil region promotes ZMM crossover recombination and is furthermore required for properly ensuring SC assembly is coordinated with (tied to) the ZMM crossover process, precisely like *zip1[Δ2-9]*, *zip1[F4A, F5A]*, and *zip1[Δ10-14]* mutants (VOELKEL-MEIMAN *et al*. 2019).

Zip3’s role in regulating the abundance of SUMOylated Ecm11 forms is also distinct from that of other ZMMs. Zip1, Zip2, and Zip4 have been found to promote the formation of SUMOylated Ecm11 in the meiotic cell, whereas Zip3 limits the formation of and/or the stability of hyperSUMOylated forms of Ecm11 (HUMPHRYES *et al*. 2013; Figure 7C, D). We found that only two of the nine *zip3* mutants examined in our study, *zip3[F190A]* and *zip3[Δ347-359]*, exhibit a wild-type distribution of SUMOylated and unSUMOylated Ecm11, while the remaining *zip3* mutants show abnormally high levels of hyperSUMOylated Ecm11 reminiscent of the *zip3* null (Figure 7C, D).

Is there a correlation between Zip3’s pro-crossover or SC assembly functions and its capacity to regulate the abundance of Ecm11 SUMOylated forms? Strictly speaking, no (Figure 7E): While five of the six alleles with a *zip3* null-like abundance of SUMOylated Ecm11 also exhibit a null-like crossover defect, the crossover defect of the *zip3[Δ96-109]* strain is less severe (Table 1). On the other hand, no correlation exists at all between the SC assembly and “SUMO-Ecm11” phenotypes - among the six *zip3* alleles with null-like abundance of SUMOylated Ecm11, some have null-like SC levels, others have less SC and still others have more SC than the null.

### Zip1 constrains Zip3 post-translational modification to be dependent on ZMM activity and Zip3 phenylalanine 190

We previously reported that post-translational modification of Zip3 during meiotic prophase relies on ZMM proteins such as Zip2 and Msh4: In *ndt80*-arrested late meiotic prophase cells missing such ZMM activity, Zip3 exhibits one discrete Zip3 species instead of multiple species of slightly different sizes (VOELKEL-MEIMAN *et al*. 2024). Here we extend this earlier analysis to demonstrate a role for Zip1 in the mechanism that links Zip3 post-translational modification to ZMM activity. We find that Zip1 removal from either the *zip2* or *msh4* mutant causes the appearance of multiple slower-migrating Zip3 forms (Figure 8A). Thus, Zip1 is essential for a mechanism that links Zip3 post-translational modification to the successful completion of ZMM activity: If the full complement of ZMMs is present in the mid-meiotic prophase cell then both unmodified and modified forms of Zip3 exist, but if a particular ZMM such as Msh4 is missing, the presence of Zip1 ensures Zip3 remains unmodified.

**Figure 8.**
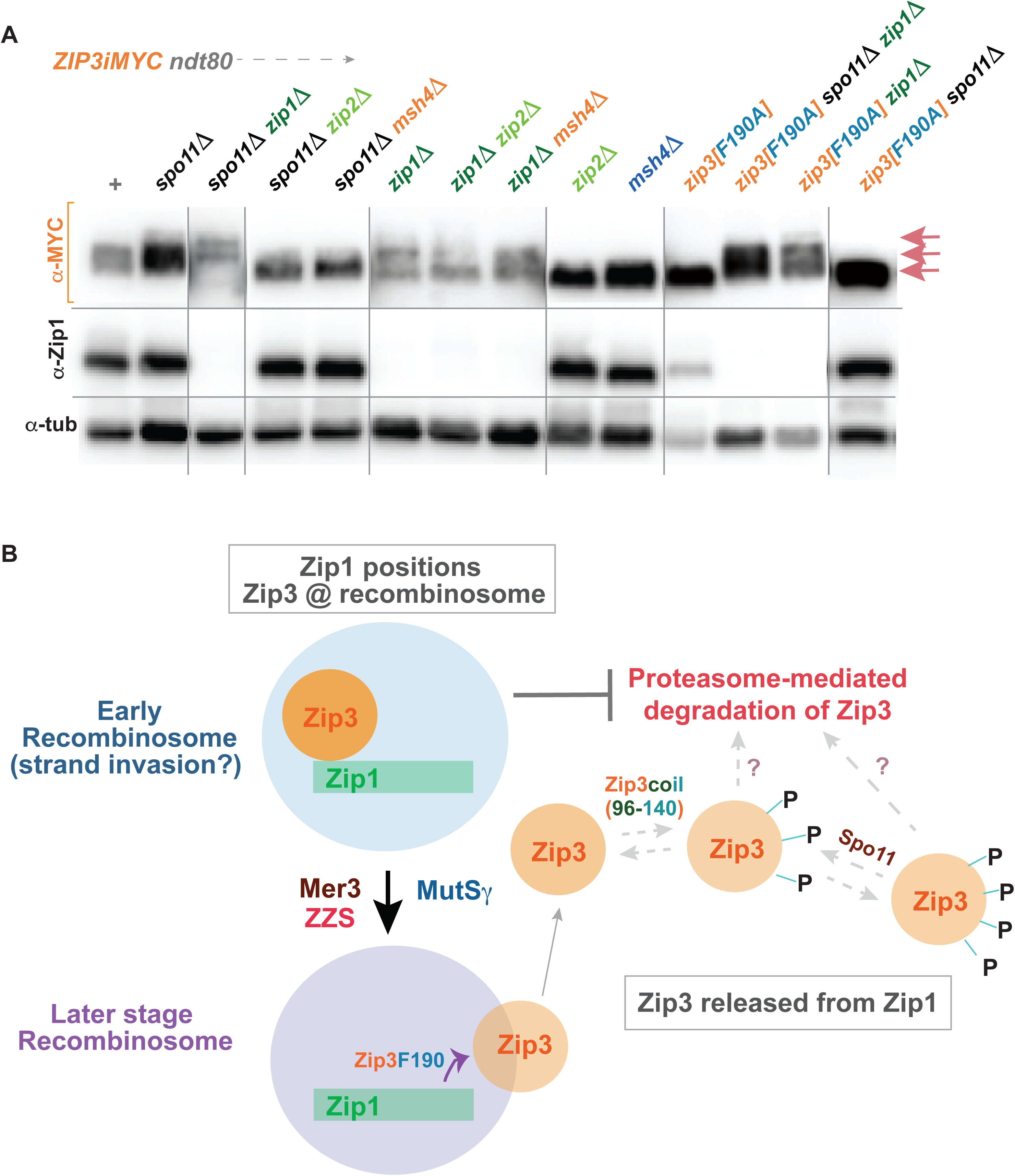
Zip1 is essential for the mechanism that links Zip3 post-translational modification to ZMM activity. Blots in (A) show proteins extracted from mid-meiotic prophase cells of *ZIP3iMYC ndt80Δ* strains missing an additional meiotic factor (listed across top of blot), separated on an 8% polyacrylamide gel. Blots were probed sequentially for anti-MYC, anti-tubulin, and anti-Zip1 antibodies. All samples are from the same blot, vertical lines indicate images from different exposure times (necessary to visualize all Zip3iMYC species in certain mutants, such as *zip1Δ*, that have low Zip3iMYC abundance). Pink arrows indicate Zip3iMYC proteins, which comprise several species of distinct sizes depending on the presence of post-translational modifications (PTMs). Blot is representative of four experiments comprising two technical and two biological replicates. (B) illustrates a model to account for the observation that, when Zip1 is present in the cell, Zip3’s capacity to transition from an unmodified to a modified form relies on ZMM activity and Zip3’s phenylalanine 190 but, when Zip1 is absent, Zip3 can undergo post-translational modification despite the absence of a ZMM or Zip3’s phenylalanine 190 (A), and furthermore that Zip3’s phenylalanine 190 is dispensable for crossover formation (Table 1). The model posits that unmodified or under-modified Zip3 is functional and stabilized by Zip1 at early recombination protein ensembles. Successful completion of intermediate steps in recombination, mediated by ZMMs, leads to Zip3’s release from Zip1, mediated by Zip3’s phenylalanine 190, and a consequent capacity of Zip3 to be post-translationally modified (likely by phosphorylation; SERRENTINO *et al*. 2013); Zip3 modification may potentially lead to an increased likelihood of degradation. Finally, the appearance of even slower migrating forms of Zip3 when Spo11 is removed from *ZIP1+* or *zip1Δ* strains (A) indicates that Spo11 activity counters, to some extent, Zip3 post-translational modification.

Removal of Spo11 from wild type or the *zip1* mutant causes additional slower migrating forms of Zip3 to appear, but this is not the case (at least not to the same extent) when Spo11 is removed from the *zip2* or *msh4* mutant (Figure 8).

Interestingly, we also found that the Zip1-dependent mechanism that ties Zip3 posttranslational modification to ZMM activity involves Zip3’s phenylalanine 190. The *zip3[F190A]* mutant exhibits nearly wild-type levels of MutSγ crossovers (Table 1). Yet, unexpectedly, *zip3[F190A] ndt80* mutants exhibit a single, fast migrating Zip3 species on a western blot like the *zip2* or *msh4* mutant. Also reminiscent of the *zip2* or *msh4* mutant, Zip1 removal from *zip3[F190A]* cells causes slower migrating forms of Zip3 to appear (Figure 8). Moreover, because the abundance of Zip3[F190A] protein is less impacted by Zip1 removal (Supplemental Figure S5) the shift in Zip3 size distribution upon Zip1 removal is more dramatic in the *zip3[F190A] zip1* double relative to the *zip2 zip1* or *msh4 zip1* double mutant. These results indicate that Zip3’s phenylalanine 190 is dispensable for MutSγ crossover formation but essential to the Zip1-mediated mechanism that ties Zip3’s posttranslational modification to the formation of these crossovers (Figure 8B).

Our analysis incidentally identified three additional disruptions to Zip3 that prevent its post-translational modification (as indicated by a single, fast migrating species on a blot; Supplemental Figure S5Bi, ii). These disruptions: *[Δ96-109]*, *[“4LE” = L109E, L119E, L127E, L140E]*, and *[Δ347-359]* are different from [*F190A]*, however, in that each negatively impacts MutSγ crossover formation (Table 1). The absence of post-translational modification of these mutant Zip3 proteins could thus be an indirect result of a failure at some intermediate step in the formation of MutSγ crossovers. This may be the case for Zip3[Δ347-359], but we find that the Zip3[Δ96-109] and Zip3[4LE] proteins are distinct from Zip3[Δ347-359] and all other altered Zip3 proteins in another way: Slower migrating forms of Zip3[Δ96-109] or Zip3[4LE] fail to appear even upon removal of Zip1 (Supplemental Figure S5Bii). Thus, the 96-140 region controls Zip3’s intrinsic capacity to be post-translationally modified during meiosis, perhaps by mediating an interaction with the post-translational modification (kinase) machinery or by harboring the residues that become modified (Figure 8B).

## Discussion

Here we integrate a new structure-function analysis of the meiosis-specific Zip3 protein with similar data for the SC component, Zip1, to gain insight into the molecular relationship between the two pro-crossover factors. Our analysis reveals several new findings about the physical nature of the Zip1-Zip3 ensemble: First, Zip3’s N terminus begins ‘MPDSIF’ (∼41 codons downstream of the translational start site currently annotated in *SGD*). Second, while residues throughout the length of Zip3 are required for normal levels of ZMM crossovers and SC assembly, the N terminal third (the only structured region of the protein) is sufficient for interacting with Zip1 *in vivo* and directly interacts with Zip1’s N terminal region in bacterial pulldown experiments. Third, disruption of a coiled region downstream of Zip3’s RING domain and within the structured N-terminal third of the protein, results in a phenocopy of the *zip1[F4A,F5A]* mutant: abolished ZMM crossovers but extensive SC assembly.

This last result aligns with the possibility that phenylalanine residues within Zip1’s N terminal tip and part of Zip3’s RING-adjacent coil act collaboratively to promote crossover recombination and prevent unscheduled SC assembly, ensuring SC formation is properly coupled with intermediate steps in the crossover pathway. As illustrated in Figure 9, we imagine this collaboration may involve the generation of a physical interface between the N-terminal region of Zip1 containing phenylalanines F4 and F5, and Zip3’s residues 81-95, which also contains a conserved phenylalanine residue (F90). Interestingly, Neves et al. (2025) recently reported that the *C. elegans* SC structural component, SYP-4, recruits a Zip3 homolog (ZHP-4) via a mechanism involving multiple phenylalanines in a C-terminal region of SYP-4 dispensable for SC formation (KOHLER *et al*. 2025; NEVES *et al*. 2025). Thus, patches of aromatic residues within disordered terminal regions of one or more SC structural proteins (which may, as in the case for SYP-4, flexibly extend outside of the more organized part of the SC lattice) may reflect a conserved structural mechanism to productively engage building block components of the SC with the factors that process recombination intermediates.

**Figure 9.**
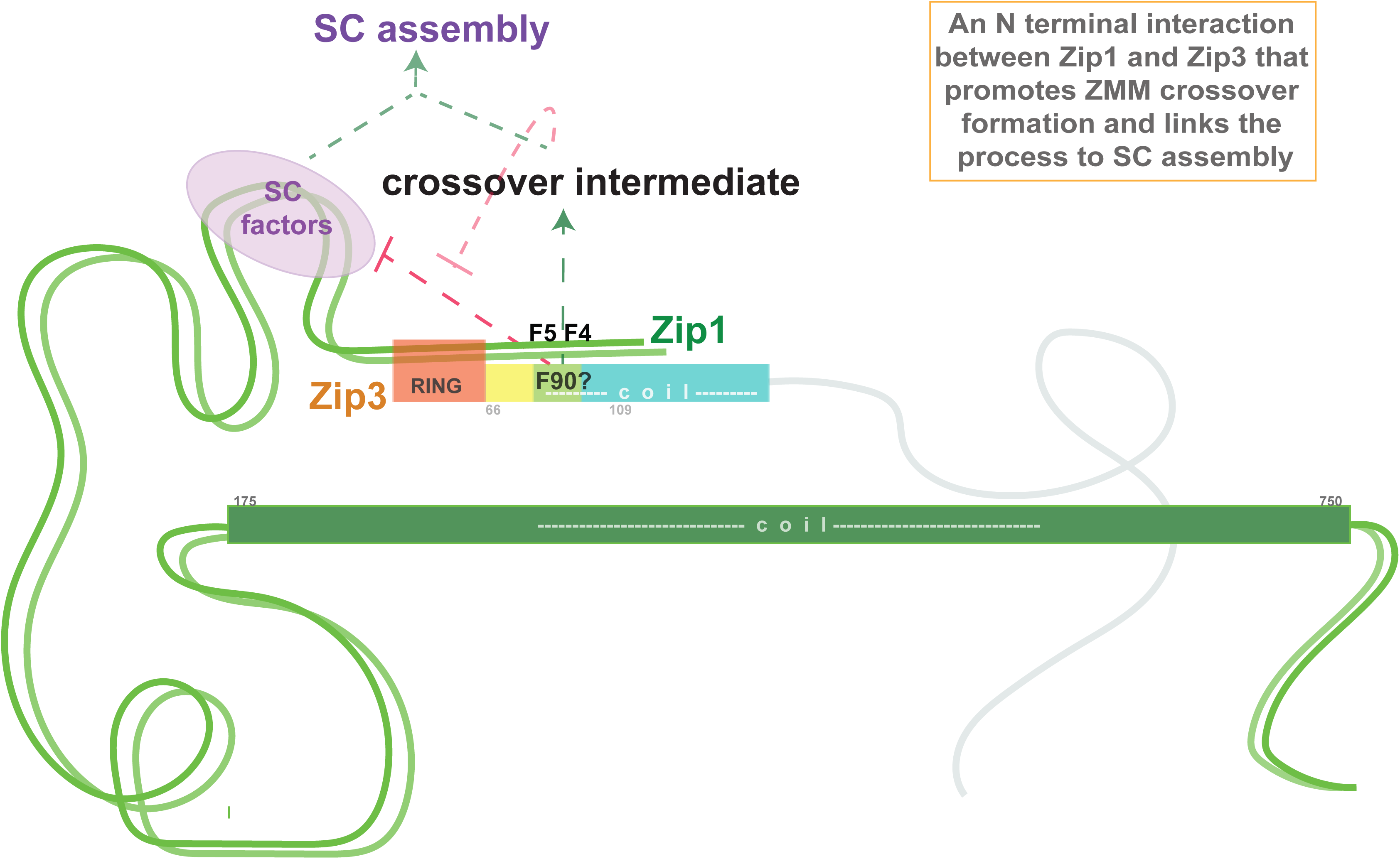
Zip1’s N terminal tip residues may interact with Zip3’s RING-adjacent coil to promote crossovers and couple SC assembly to the crossover process. Cartoon illustrates Zip1 and Zip3 proteins, with N-terminal residues of each forming an interface, particularly between Zip1’s F4, F5 region and Zip3’s residues 81-95 (lime green). The illustration highlights a conserved phenylalanine, F90, in this 81-95 region of Zip3. This interface is proposed based on phenotypic similarity between *zip3[Δ81-95]* and *zip1[F4A, F5A]* or *zip1[Δ2-9]* mutants (VOELKEL-MEIMAN *et al*. 2019; VOELKEL-MEIMAN *et al*. 2024; this work) and we speculate the interaction facilitates crossover formation and prevents unscheduled SC assembly: SC formation in the absence of ZMM crossovers (VOELKEL-MEIMAN *et al*. 2019). Residues downstream of this pro-crossover of Zip1’s N terminus are essential for SC assembly (VOELKEL-MEIMAN *et al*. 2016) through a mechanism coordinated with intermediate steps in recombination, as depicted in the flowchart.

Curiously, meiosis-specific RING proteins like Zip3 are conserved players of the meiotic recombination pathway (AGARWAL AND ROEDER 2000; TOBY *et al*. 2003; JANTSCH *et al*. 2004; KONG *et al*. 2008; STRONG AND SCHIMENTI 2010; CHELYSHEVA *et al*. 2012; REYNOLDS *et al*. 2013; QIAO *et al*. 2014; LAKE *et al*. 2015; LAKE *et al*. 2019), but it remains unclear why; what conserved function(s) do they provide? While *S. cerevisiae* Zip3 is capable of E3 SUMO ligase activity *in vitro* (CHENG *et al*. 2006), how this activity relates to its critical pro-crossover or SC regulatory activity *in vivo* is not understood. When Zip3 is absent, SUMOylated forms of Ecm11 increase in abundance about 1.5-2-fold (Figure S2C), consistent with the idea that Zip3 either prevents and/or attenuates the formation of SUMOylated Ecm11, or perhaps acts as a STUbL (SUMO-Targeted Ubiquitin Ligase; GARZA AND PILLUS 2013) to promote the turnover of SUMOylated Ecm11. Another possibility, which is not mutually exclusive, is that Zip3 acts not only in SUMO or ubiquitin enzymatic pathways but also serves a structural role as a binding partner for other proteins and/or DNA within the recombinosome ensemble. The data presented here highlight not only that Zip3’s structured N terminus is necessary and sufficient for interacting with the N-terminal region of SC structural component Zip1, but also that the unstructured C terminal end of the polypeptide plays a critical role in Zip3’s crossover and SC regulatory activities. Identification of factors interfacing with this extensive unstructured region of Zip3 may shed light on how this RING protein facilitates successful meiotic recombination.

The results presented here indicate that Zip1’s direct interaction with Zip3 is one critical mechanistic aspect of the ZMM crossover pathway, and that this interaction protects Zip3 not only from proteasome-mediated degradation but also from post-translational modification in the context of an incomplete or non-functional ZMM cohort. Considering our previously published observation that Zip3 post-translational modification depends on a full cohort of ZMM activity (VOELKEL-MEIMAN *et al*. 2024), and that Zip3[F190A] remains unmodified (Figure 8) during otherwise-normal meiosis but is functional as a crossover protein (Table 1), we propose that Zip3’s interaction with Zip1 at a recombination intermediate may serve in a checkpoint-like manner, to ensure Zip3 remains in an unmodified state within the context of an incomplete recombinosome ensemble, until a full cohort of ZMM factors and/or a mature joint molecule DNA recombination intermediate is present.

## Data Availability

Raw data presented in graphs or used to calculate average measurements are provided in Supplemental File S7.

## Acknowledgements

We are grateful to MB&B 394 Advanced Lab students: Erica Horowitz for generating multiple *zip3* alleles, Phyllis Schram and Aiden Parente for creating the *ZIP3* expression plasmid and carrying out initial co-expression and pull-down experiments. We also thank Dr. Owen Davies for supplying the pMAT11-Zip1 expression plasmid, and Dr. Andrea Pichler for suggesting the idea that Zip3 functions as a STUbL. We thank Drs. Scott Holmes and Michael Weir for valuable feedback on experiments and the manuscript. The work was supported by National Institutes of Health grant GM116109 to A.J.M.

